# Quantification of Local Mesostructural Deformation of the Myocardium using DENSE MRI and a Novel Set of Strain Attributes

**DOI:** 10.64898/2026.11.11.766054

**Authors:** Tawfik M. Hussein, John N. Oshinski

## Abstract

Measuring myocardial deformation is central to understanding the mechanical behavior of both the healthy and the diseased heart. While myocardial deformation is quantified via medical imaging in the form of strain, numerous definitions have been used in the engineering and clinical literature, resulting in inconsistencies in interpreting mechanistic changes in myocardial function. Additionally, current strain definitions lack the robustness required to relate quantitative changes in strain to changes in the myocardial mesostructure. Hence, the goal of this work was to propose a novel set of logarithmic, mesostructure-based strain attributes, by leveraging the upper triangular (QR) decomposition theorem. These strain attributes were then compared with the polar Green strain, the most widely used strain definition for the myocardium. Ten healthy subjects were scanned via displacement encoding with stimulated echoes (DENSE). From these images, a 3-dimensional (3D) displacement field was obtained, from which both 3D Green strains and strain attributes were calculated and compared. The results of this work revealed that the two strain definitions yield significantly different deformation profiles of the myocardium (p < 0.05), with strain attributes predicting relatively more spatial heterogeneity compared to the Green strains — a finding consistent with similar studies that quantified mesostructural myocardial deformation. Additionally, the strain attributes revealed the myocardium to be compressible, challenging the common assumption of incompressibility. By capturing the deformation along the mesostructure, the strain attributes enable intuitive characterizations of cardiac disease, providing clinicians with deeper mechanistic insights into left ventricular function and serving as predictive markers of adverse remodeling.

## INTRODUCTION

Quantifying myocardial deformation is central to understanding the physiological behavior of the healthy heart as well as interpreting pathophysiological phenomena in the diseased heart [1–8]. Measuring the deformation of the myocardium carries many advantages as it is defined locally and is specific to myocardium itself, unlike global functional measurements such as ejection fraction (EF). EF can be less sensitive to regional changes within the myocardium and can be driven by changes in cardiac structures other than the myocardium. Therefore, myocardial deformation can be used as a robust clinical marker of myocardial function in various cardiac diseases, as a metric for evaluating the efficacy of cardiac devices and therapies, or as a method to quantify how well animal disease models represent the cardiac disease of interest [1–4,9–16]. Myocardial deformation can also be used to computationally simulate the mechanical behavior of digital twins of patient-specific hearts [17–19]. While myocardial deformation is quantified as strain, various definitions of strain have been used, resulting in differences in how deformation has been interpreted in various disease states. For instance, the engineering strain and the Green strain have both been widely used as definitions for myocardial strain. However, these definitions have numerous limitations (as discussed in the following section), so the primary purpose of this work was to establish a strain measure that can robustly capture myocardial deformation and can be directly connected to physiological changes. This proposed strain measure, the strain attributes, employs a mesostructural description of myocardial deformation. The goal of this work was two-fold: (1) quantify these mesostructure-based strain attributes in the left ventricle (LV) of healthy human subjects; and (2) compare the strain attributes to the polar Green strains for capturing changes in the myocardial mesostructural architecture. Because we had chosen the region of interest to be over the full transmural thickness of the myocardium, we assumed the mean fiber direction to be aligned with the hoop direction. As a result, the material-based coordinate system herein is aligned with the polar axes. This granted us the unique opportunity to test our hypothesis that the strain attributes will yield a deformation that is significantly different from that of Green strain, with greater spatial heterogeneity that is consistent with myocardial physiology and changes in myocardial architecture with contraction. We also compared the strain attributes with engineering strain to illustrate that engineering strain is not appropriate to use for the myocardium.

## BACKGROUND & THEORY

### Myocardial Deformation – A Consequence of the Myocardial Architecture

The specific mechanism of cardiac deformation arises from the organization of the myocytes within the myocardium as well as the collagenous perimysium occupying the extracellular matrix [20–23]. The myocardium of the left ventricle (LV) has a complex architecture, such that the myocardial fibers are combined into myocardial ‘sheetlets’ that are roughly 4 cells thick, and these sheetlets are separated from one another via cleavage planes [5–8,15,20–24]. These sheetlets are ‘stacked’ through the myocardium transmurally from the endocardium to epicardium, with each sheetlet having a distinct orientation [5–8,15,20–24]. The myofibers within each sheetlet is oriented at an angle from the LV equator, with a transmural gradient from endocardium to epicardium, where the subendocardial fibers subtend an angle of ∼60° with the equatorial axis; the mid-myocardial fibers subtend a 0° angle (i.e., aligned with the hoop direction); and the subepicardial fibers subtend a ∼ – 60° angle (demonstrated in **Figure 1**) [15]. This angle is known as the ‘in-plane’ angle, which is the angle of the fibers within the plane of the sheetlets. There are two other angles important to myocardial structure related to the amount of sheetlet imbrication: one resulting from sheetlet tilting about the equatorial axis, and the other resulting from sheetlet tilting about the longitudinal axis [5–8,15,20–23]. Both of these angles are known as the two ‘out-of-plane’ angles as they are related to the angle between the sheetlet outward normal and the local radial axis and the circumferential axis, respectively. Hence, when the myofibers contract, the architectural organization of the sheetlets produces a global deformation pattern that results in LV myocardial thickening by ∼40% (due to the increased shearing of the imbricated sheetlets at systole) and shortening along the myocardial circumferential and longitudinal axes [5–8,15,20–23].

**Fig. 1.**
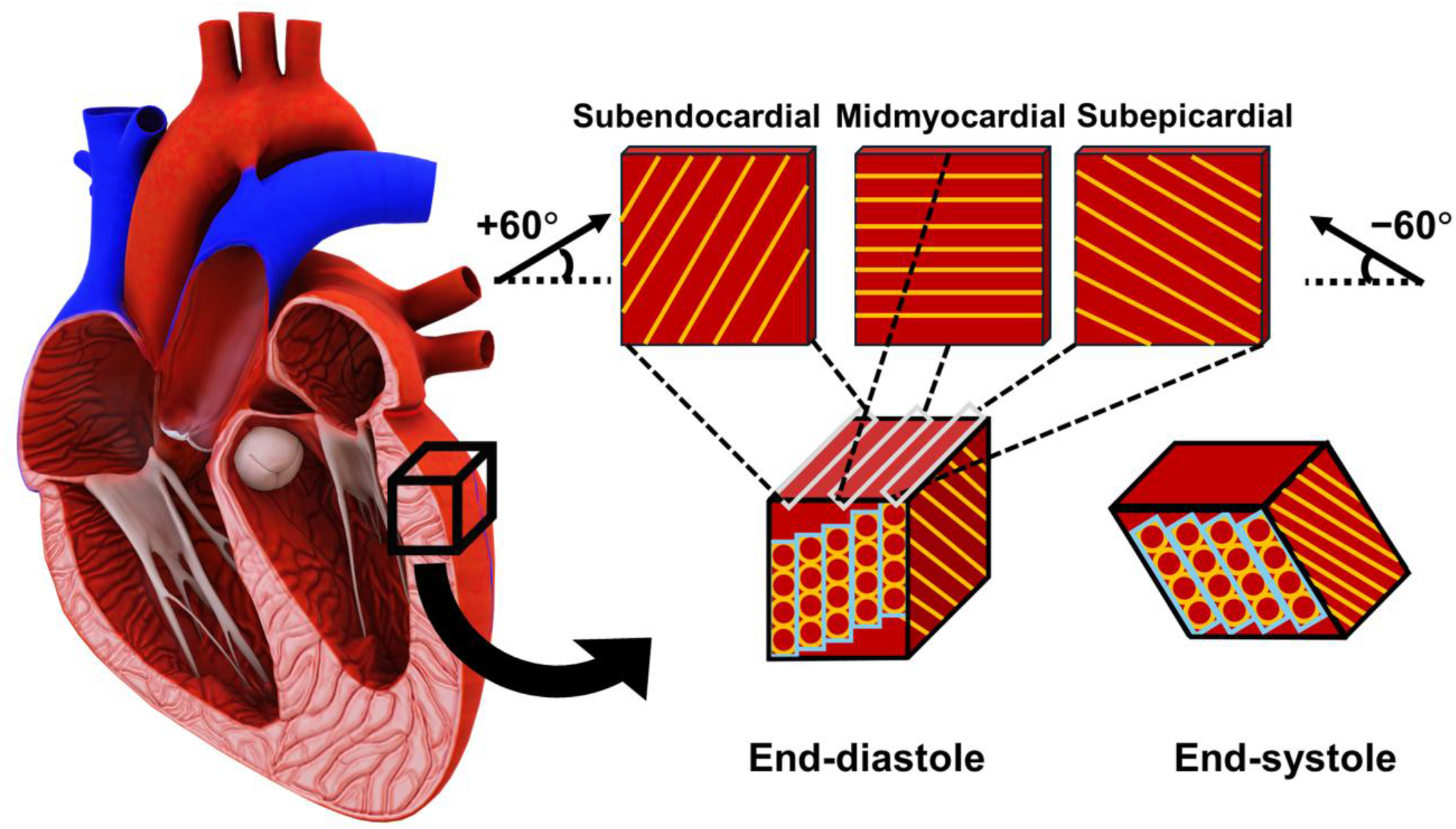
Architectural organization of the left ventricular myocardium, where the fibers are shown from subepicardium to the subendocardium, with the myocardial-sheetlet shearing shown in end-diastole and end-systole.

[Figure 1 HERE]

### Myocardial Strain – A Quantification of the Myocardial Deformation

Broadly, strain is a tensor that quantifies the amount of displacement between two material points along any given direction. It is thus locally defined and described by a 3 × 3 matrix that is usually symmetric. Myocardial strain can be estimated *in vivo* non-invasively from echocardiography, computed tomography (CT), and magnetic resonance imaging (MRI) [2,4,11,25–27]. With MRI, strain can be quantified via feature-tracking (FT) based on cine balanced steady state with free precession (bSSFP) images, myocardial tissue tagging, phase contrast tissue velocity mapping, or displacement encoding with stimulated echoes (DENSE) [1–4,11,28–32]. Due to its high spatial resolution in directly quantifying a pixelwise displacement field, DENSE MRI is considered the gold standard in measuring myocardial strain [1–4,33–38].

There are 2 main definitions of strain that are used across the clinical and engineering communities: the engineering strain and the Green strain, respectively. The engineering strain is defined as the symmetrical part of the displacement gradient tensor (i.e., **ε** = *sym* [**Grad(u)**], where **ε** is the engineering strain tensor; **u** is the displacement field; and **Grad()** is the material gradient operator); however, what is often used is the one-dimensional (1D) definition of engineering strain, i.e., *ε*_11_ = Δ*L*/*L_0_*, where *L_0_* is the 1D reference length (typically taken at end-diastole), and Δ*L* = *L – L*_0_, where *L* is the current or deformed length [1–4,11,14]. While 1-D engineering strain is commonly used in clinical studies and implemented by several commercial strain analysis packages, the engineering strain simply does not apply to the myocardium (or any other soft tissue) [1,3,4,11,26,39]. Engineering strain only applies to materials that undergo “small” or infinitesimal deformations, usually defined at less than 2% elongation or shortening such as metals, alloys, and wood which fall within the realm of linear elasticity [40]. However, the myocardium undergoes “large” or finite deformations, often exceeding stretches of 30% and thus falling within the realm of non-linear elasticity [41]. Employing engineering strain for the myocardium is, therefore, not appropriate.

The Green strain, on the other hand, is a definition of strain that is suitable for non-linear materials undergoing large deformations (defined as 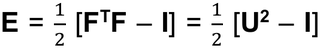, where **E** is the Green strain tensor; **F** is the local deformation gradient tensor; **F^T^** is the transpose of **F**; **I** is the identity tensor; and **U** is the right Cauchy-Green stretch tensor). The Green strain has been previously quantified via MRI in humans and large animals, where the Green strain was measured along polar axes or global cardiac coordinates (i.e., circumferential, longitudinal, and radial) which define the deformation in terms of changes to the myocardial geometry [11,15,16,30,42,43]. Furthermore, circumferential Green strain (*E*_CC_) defines the amount of shortening of the myocardial circumference; radial Green strain (*E*_RR_) defines the amount of radial thickening of the myocardium; and longitudinal Green strain (*E*_LL_) defines myocardial shortening along its longitudinal axis. The components are shown in the tensor expression below:

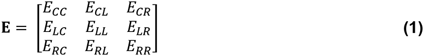

The off-diagonal components define shear distortions of the relevant planes, whereby E_CL_ is the shearing of the circumferential-longitudinal plane (often related to myocardial twist); E_CR_ is the shearing of the circumferential-radial plane (which arises due to the tilting of the sheetlets about their longitudinal axis); and E_LR_ is the shearing of the longitudinal-radial plane (which arises due to the tilting of the sheetlets about their circumferential axis). These components can be visualized from **Figure 2**. Hence, for any strain tensor, the diagonal components quantify the *normal strains* (i.e., deformation along the axes) whereas the off-diagonal components quantify the *shear strains* (i.e., deformation due to shearing of the planes).

**Fig. 2.**
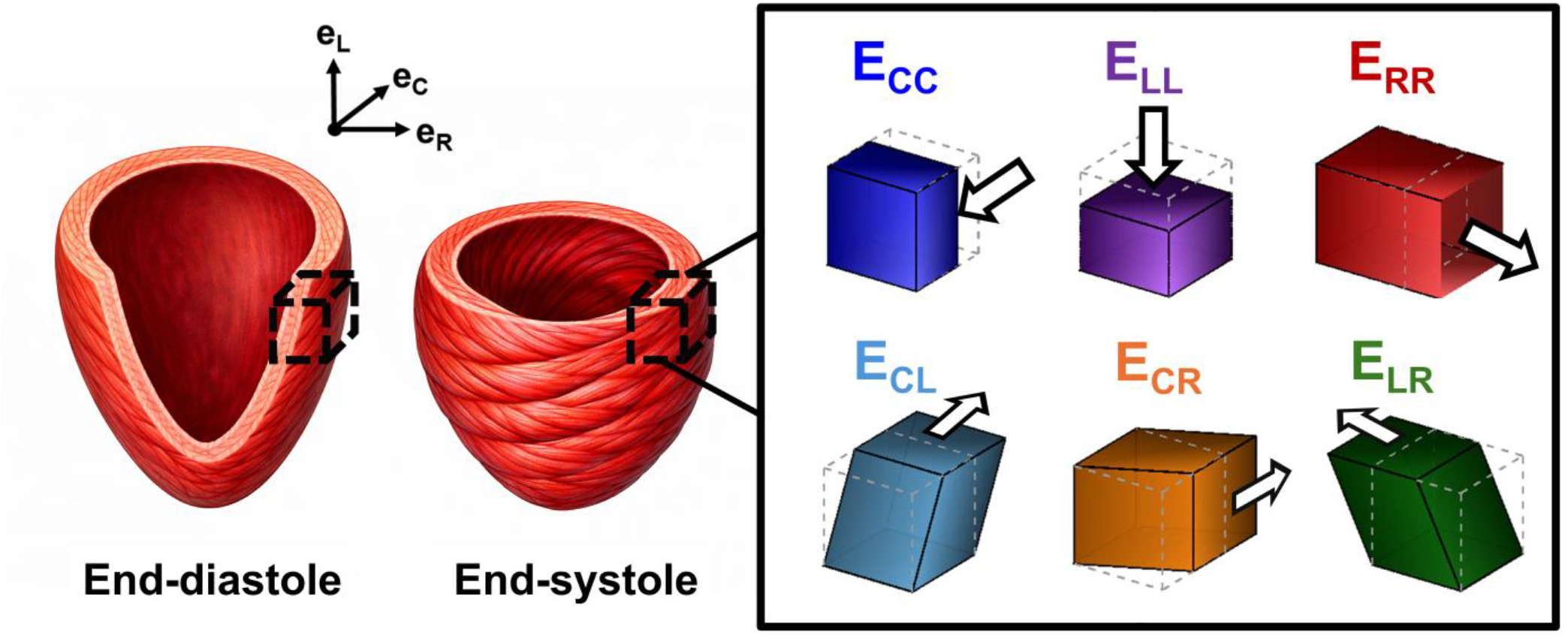
Myocardial modes of deformation per the components of the Green strain tensor demonstrated on a cube taken from the base of the left ventricle.

[Figure 2 HERE]

### Limitations of the Green Strain

While the polar Green strains (also regarded as *geometric* Green strains) are suitable for quantifying the large deformations along the polar geometry of the myocardium, they suffer from two main limitations: they are neither physical (i.e., not as physiologically intuitive) nor do they *directly* describe changes in the myocardial mesostructure. Regarding its physicality, the Green strain is a quadratic function of the stretch, where the stretch is squared, subtracted by unity, and then halved (e.g., 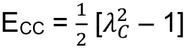). Hence, it is not indicative of the ‘true’ amount of elongation or shortening experienced by the myocardium. Additionally, regional distributions of the polar Green strain from MRI show a homogeneous distribution of the circumferential strain, longitudinal strain, and radial strain, where they are roughly equal in magnitude but are opposite in signs (radial is positive while both circumferential and longitudinal are negative).

However, investigations that studied myocardial deformation along microstructurally defined directions (i.e., fiber, sheet, and normal) show much greater regional heterogeneity [5–8,15,16]. Because polar Green strains merely describe the geometric changes undergoing the myocardium, quantifying deformations along the mesostructure is more congruent with changes in myocardial architecture and physiology, which can better describe pathophysiological phenomena [34,44]. It is important to note herein that the term *mesostructure* refers to length scales in the order of millimeters (e.g., myofibers and sheetlets), which is distinct from the term *microstructure* that refers to length scales on the order of microns (e.g. cardiomyocytes and subcellular structures). Regarding the mesostructure, because polar Green strains merely describe the geometric changes undergoing the myocardium, quantifying deformations along the mesostructure has shown to be more congruent with myocardial physiology [34,44]. Thus, capturing myocardial deformation along mesostructurally relevant directions can reveal changes in the myocardial architecture and can better describe pathophysiological phenomena [34,44]. This would potentially make mesostructure-based definitions of strain more clinically intuitive and physically relevant. Additionally, a mesostructural description of myocardial deformation is yieldable by DENSE MRI because the length scales of the mesostructure is consistent with the length scales obtainable by the voxel size of DENSE.

### The Strain Attributes – A Mesostructural Description of Myocardial Deformation

Recently, Srinivasa developed a theory to describe material deformation relative to the mean material axes (as opposed to basis coordinates), which yields a decomposition of the local deformation gradient tensor **F** into an upper triangular stretch tensor **f** and a pure rotation tensor **Q**, known as the upper triangular decomposition (i.e., **F** = **Qf**) [45]. The upper triangular stretch

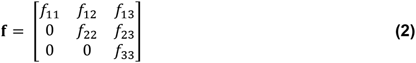

This theory has been shown to result in stretches that portray physical deformations along the material axes in addition to other important advantages outlined below [45]. For the myocardium, these axes correspond to mesostructurally-defined axes, which are the *mean* fiber axis, cross-fiber axis (in-plane perpendicular), and radial axis (out-of-plane perpendicular), where the mean is taken over the region of interest. Hence, the components of the upper triangular stretch tensor shown in **Eq. 2** describe myocardial deformation as follows: (1) *f*_11_ is the stretch along the mean fiber direction; (2) *f*_22_ is the stretch along the cross-fiber direction; (3) *f*_33_ is the stretch along the radial direction; (4) *f*_12_ is the stretch resulting from shearing the cross-fiber face along the mean fiber axis; (5) *f*_13_ is the stretch resulting from shearing the fiber face along the radial direction; and (6) *f*_23_ is the stretch resulting from shearing the cross-fiber face along the radial direction. To clarify, stretch (*λ*) is defined in 1D as the ratio of the current length (*L*) to the initial length (*L*_0_), or *λ* = *L*/*L*_0_. For example, if a slab was elongated by 40% of its original length, it is said to have undergone a stretch of 1.4 (whereby *L*/*L*_0_ = [*L*_0_ + 0.4*L*_0_] / *L*_0_ = 1.4).

From these upper triangular stretches, Criscione et al. proposed a set of strain attributes that can be derived directly from these stretches that are optimal for laminar materials with one-fiber family, such as the myocardium [24,41]. Rather than being related to the stretches quadratically (such as the Green strain), these strain attributes are *logarithmic* functions of the stretch, resulting in deformations that are more physical and robust. Furthermore, logarithmic strain results from the sum of the infinitesimal subregions of a finitely deformed region along the direction of interest [46–50]. This is shown below in **Eq. 3**, where *ξ* is the logarithmic strain, *x* is the deformed region in the current configuration, *dx* is an infinitesimal neighborhood within the deformed region in the current configuration, and λ is the stretch ratio between the current length (*L*) and initial length (*L*_0_). **Figure 3** can be referenced for visualization, which shows a black strip with initial length *L*_0_ and red region Δ*X*. When the strip is stretched, its current length becomes *L*, and its current red region becomes Δ*x*. To obtain the deformation (i.e., the change from Δ*X* to Δ*x*), the small neighborhoods or increments within Δ*x* (i.e., *dx*) are added together. To obtain the amount of deformation relative to the current length, the increments are added as ratios (i.e., *dx/x*). This yields the integral in **Eq. 3**, which results in the natural logarithm of the stretch. Additionally, logarithmic strains have shown to have other advantages, including inverse symmetry, no negative or positive bias, and allowing separability, where the deformation can be separated as a summation of dilatation and distortion (i.e, **ln [**(*dilatation*)*(distortion*)] = **ln[***dilatation*] + **ln[***distortion*]) [46–50].

**Fig. 3.**
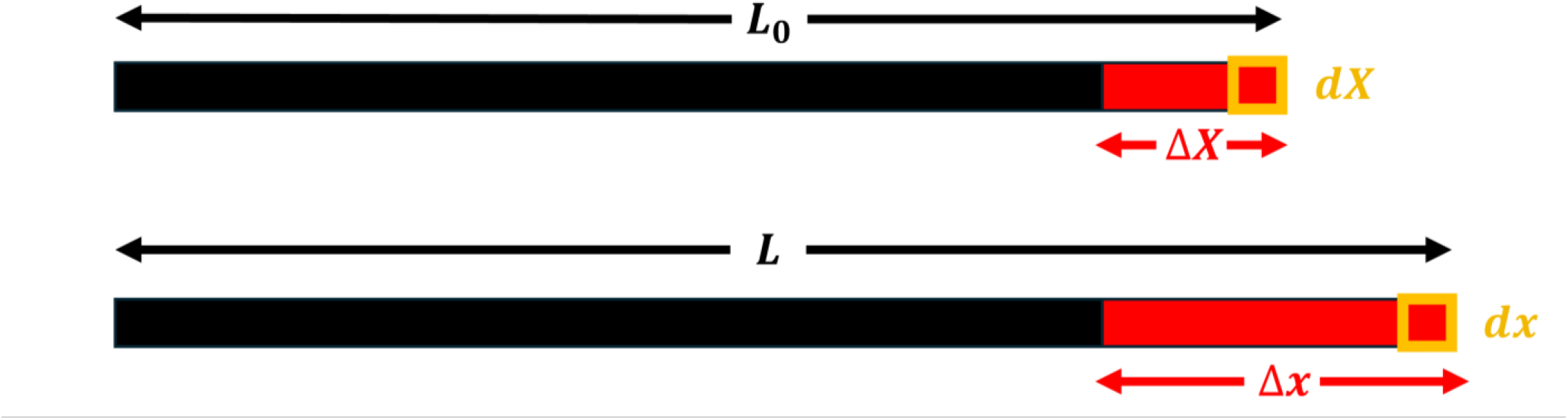
One-dimensional elongation of a strip demonstrates the definition of the logarithmic strain, which is the sum of the infinitesimal regions **dx** along the deformed region Δ**x**, per the current length **x**.

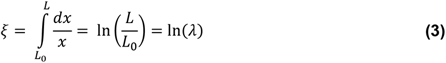

Hence, the strain attributes combine the advantages of logarithmic strain with a mesostructural description of myocardial deformation based on the upper triangular decomposition framework. Besides such advantages, the strain attributes can play a significant role in characterizing myocardial disease. Because numerous types of cardiac disease can disrupt the normal mesostructure of the myocardium, such disruptions can be captured by the strain attributes. Additionally, numerous cardiomyopathies could alter the myocardial mesostructure without necessarily altering the overall myocardial geometry during the early stages of the disease [51]. As a result, the strain attributes can help delineate the subclinical changes undergoing the myocardium, thus playing a significant role in predicting heart disease [51]. Other advantages and disadvantages of both the geometric and mesostructural descriptions of the deformation have been discussed in a previously published review [51].

## METHODS

### Participants & Imaging Protocol

Ten healthy subjects (6 males, age 25 – 59 years and 4 females, age 22 – 31 years) were recruited for this study, which was approved by the Emory Institutional Review Board (IRB). All subjects provided informed written consent and were scanned on a *MAGNETOM PrismaFit* 3T MRI scanner using a multi-element body array coil (Siemens Healthineers; Erlangen, Germany). For each subject, cine ECG-gated 2D DENSE was acquired in 6 imaging views: 3 slices along the cardiac short axis and 3 slices along the cardiac long axis [37]. The 3 short-axis slices were at the basal, mid-ventricular, and apical levels of the myocardium while the 3 long-axis slices were at the two-chamber (2CH), three-chamber (3CH), and four-chamber (4CH) views. The DENSE pulse sequence employed was a spiral DENSE acquisition protocol with outer volume suppression using the following imaging parameters: slice thickness = 8 mm, TR = 15 ms, TE = 1.08 ms, cardiac phases = 22 – 25 (depending on the subject’s heart rate), FOV = 200 mm^2^, matrix size = 128 × 128, pixel size = 1.6 × 1.6 mm^2^, 2D in-plane displacement using the simple 3-point method displacement encoding frequency = 0.1 cycles/mm, through-plane dephasing frequency = 0.08 cycles/mm, flip angle = 15°, fat suppression, and 4 spiral interleaves with 2 interleaves acquired per heartbeat [35–37,52,53]. Each view was acquired at end-expiration over a breath-hold.

### DENSE Post-processing

**Fig. 4.**
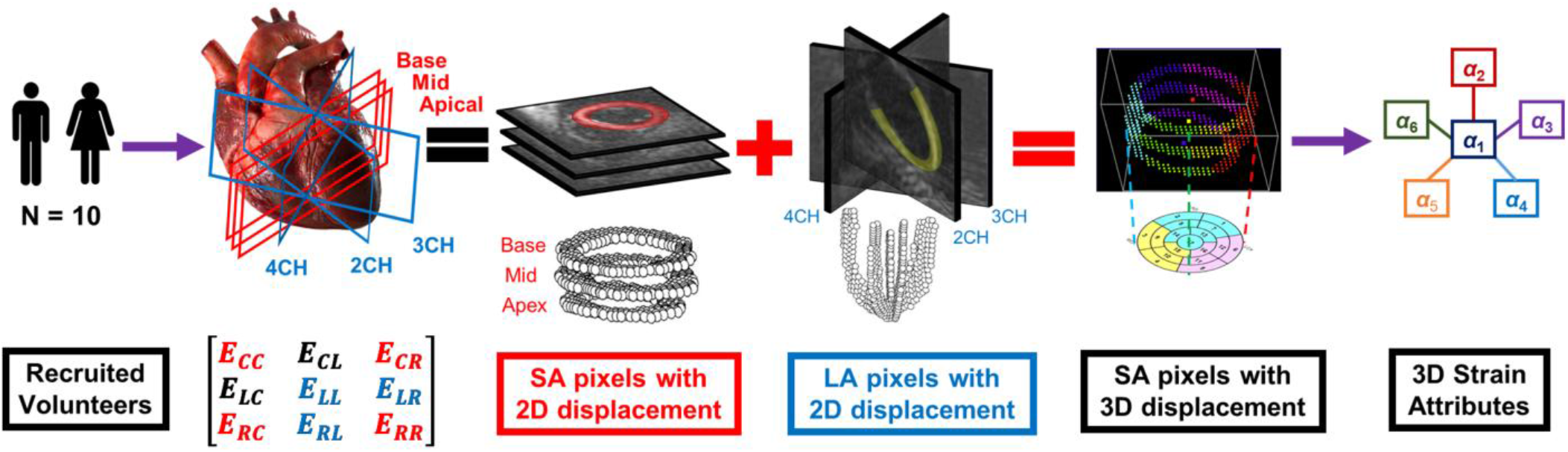
DENSE post-processing pipeline, where 10 healthy volunteers were scanned. For each volunteer, 3 short-axis and 3 long-axis images were obtained and combined to obtain an AHA 17-segment model of 3D strain and strain attributes.

The DENSE post-processing and strain analysis workflow is graphically summarized in **Figure 4**. For each of the 6 views, 3 sets of DENSE images were acquired: one magnitude image and two phase images — one along each of the Cartesian *x* and *y* motion encoding directions (in the coordinate system of the MRI scanner). Post-processing was done using the open-source *DENSEanalysis* MATLAB tool (https://github.com/denseanalysis/denseanalysis) [54]. The images were imported into *DENSEanalysis* where segmentation of the LV was done semiautomatically using motion-guided segmentation, and manual corrections were done as needed [55]. Phase-wrapped pixels were unwrapped and the Lagrangian displacement and Green strain were calculated for each pixel in the LV myocardium [56]. To obtain the polar strains, the 2D Green strain tensor obtained from the short-axis images were projected to circumferential and radial axes relative to the LV center of mass, resulting in the E_CC_ and E_RR_. Likewise, the 2D Green strain tensor obtained from the long-axis images were projected to the longitudinal and radial axes, resulting in E_LL_ and E_RR_.

Additionally, the 2D Green strain tensor was projected to the principal axes to obtain the principal Green strains in both views: the first and second principal strains obtained from the short-axis images (E_11_ and E_22_, respectively), and the first and second principal strains obtained from the long-axis images (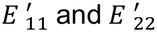, respectively). It should be worth noting that the principal axes **e_1_** and **e_2_** of the short-axis images do not form the same right-handed coordinate system with the principal axes of long-axis images. This implies that neither 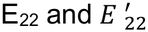 are the same nor is 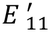 the through-plane principal strain of the short-axis E_11_ and E_22_ (i.e., 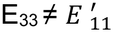).

Finally, to standardize the visual representation of the LV myocardium, both the polar and principal strains in the short-axis images were represented segmentally based on the American Heart Association (AHA) 17-segment model (also known as the AHA bullseye plot), consisting of 6 basal and 6 mid-ventricular segments as well as 4 apical segments (the 17^th^ segment, corresponding to the apex, is not presented as all 3 strain components could not be measured in that segment) [57]. The 6 segments in the base and mid correspond to the anterior, anteroseptal, inferoseptal, inferior, inferolateral, and anterolateral segments whereas the 4 segments in the apical myocardium correspond to the anterior, septal, inferior, and lateral segments [57]. For the long-axis strains, the myocardium was divided into 7 segments: 2 basal, 2 mid-ventricular, 2 apical, and 1 apex. Therefore, the final output of the DENSE post-processing step are the segmental E_CC_, E_RR_, E_11_, and E_22_ from the short-axis images, and segmental 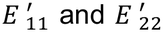 from the long-axis images (E_RR_ from the long axis was ignored since it was already obtained from the short-axis images). The segmental strains were calculated over the cardiac cycle.

### 3D Green Strain Analysis

For each subject, the DENSE post-processing step produced segmental normal strains *E*_CC_, *E*_RR_, *E*_11_, and *E*_22_ from the short-axis images and *E*_LL_, 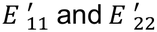 from the long-axis images. As shown in **Figure 4**, to obtain the full 3D strain tensor in **Eq. 1** from the 2D strains, four steps were needed: (1) assembling the diagonal strain tensor components from the normal strains in the short-axis segments (i.e., *E*_CC_ and *E*_RR_) and their corresponding through-plane strain in the long-axis segments (i.e., *E*_LL_); (2) using the segmental twist angle to calculate segmental E_CL_; (3) calculating E_CR_ from the short-axis polar and principal strains (*E*_CC_, *E*_RR_, *E*_11_, and *E*_22_); and (4) calculating E_LR_ from the long-axis polar and principal strains (*E*_LL_, *E*_RR_, 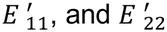).

For the first step (illustrated in **Figure 5**), the 3 ventral segments of the 2CH view corresponded to the basal anterior, mid anterior, and apical anterior segments of the short-axis whereas the 3 dorsal segments corresponded to the basal inferior, mid inferior, and apical inferior segments of the short-axis images. For the 3CH view, the 3 ventral segments corresponded to the basal anteroseptal, mid anteroseptal, and apical anteroseptal short-axis segments whereas the 3 dorsal segments corresponded to the basal inferolateral, mid inferolateral, and apical inferolateral short-axis segments. For the 4CH view, the 3 ventral segments corresponded to the basal anterolateral, mid anterolateral, and apical anterolateral short-axis segments whereas the 3 dorsal segments corresponded to the basal inferoseptal, mid inferoseptal, and apical inferoseptal short-axis segments. Hence, using this paradigm, the segmental through-plane E_LL_ strains from the long axis were locally matched with their corresponding segmental strains in the short-axis, resulting in a complete AHA 17-segment model for E_LL_.

**Fig. 5.**
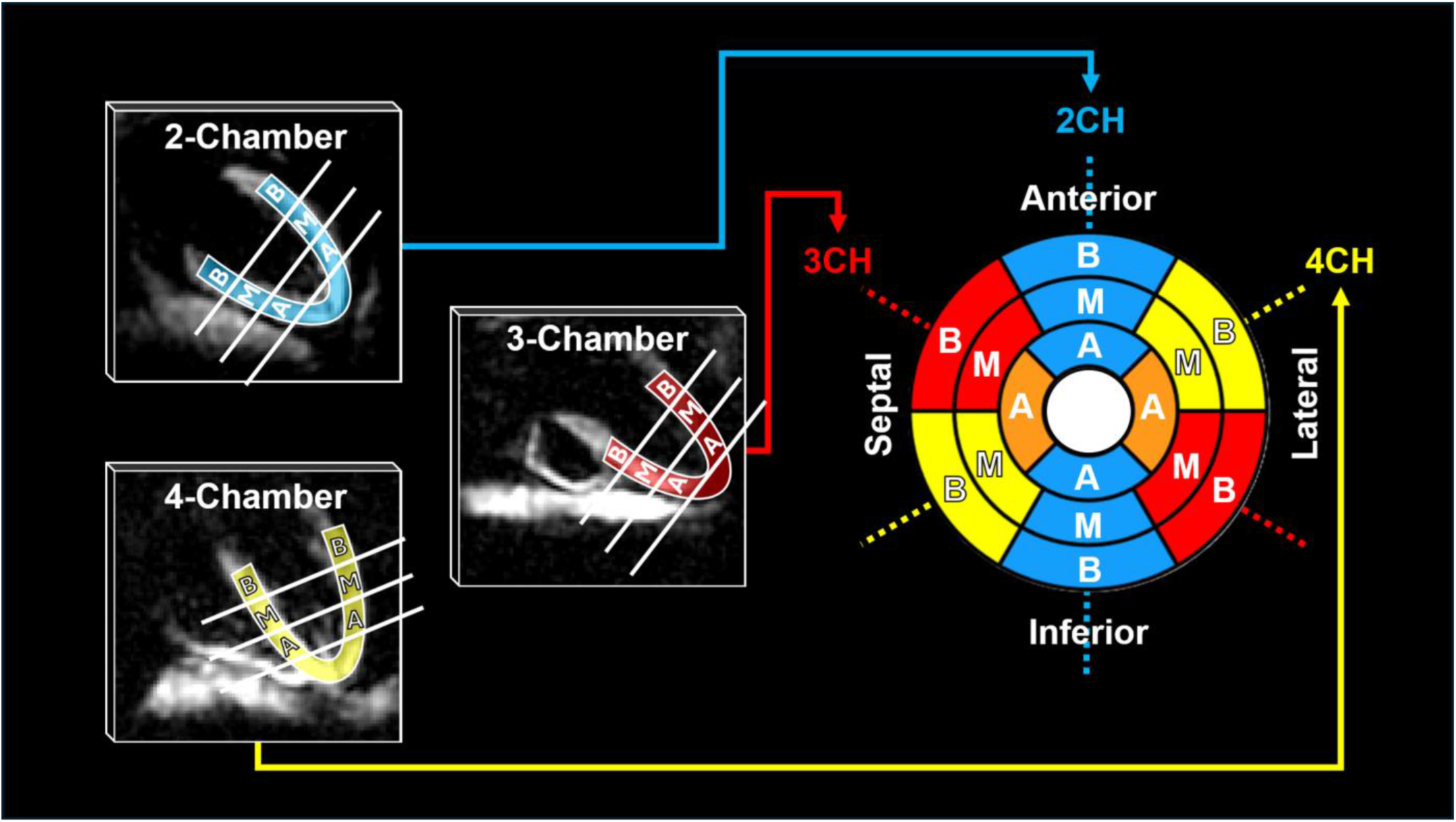
Mapping the through-plane strains from each segment in the long-axis images to their respective segment locations in the AHA bullseye. B = Base; M = Mid; A = Apex; and CH = Chamber.

For the second step, segmental E_CL_ was calculated from the segmental twist angles for each subject. Furthermore, given two short-axis slices, their relative arcual displacement per segment can be used to calculate the relative shear angle per segment via **Eq. 3a**. From the shear angle, E_CL_ can be obtained segmentally using E_CC_ and E_LL_ of the corresponding segment via **Eq. 3b**.

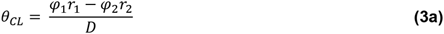

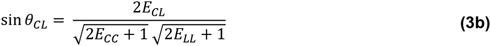

where *θ*_CL_ is the shear angle; *φ*_1_ and r_1_ are the segmental twist angle and radius of the upper short-axis slice; *φ*_2_ and r_2_ are the segmental twist angle and radius of the lower short-axis slice; and *D* is the distance between both slices. It is important to note that the *absolute* arcual displacement for each short-axis slice was of interest rather than the *relative* arcual displacement to obtain the absolute E_CL_ for the basal, mid, and apical slices. From previous studies of myocardial torsion (i.e., twist per unit length), the base twists clockwise (represented conventionally with a negative sign) whereas the apex twists counterclockwise (represented conventionally with a positive sign) with an increasing gradient of twist from base to apex, when viewed from the apex [58]. Hence, it was assumed in this paper that a slice exists where there is no twist (between the base and mid), which was used to find the absolute twist and shear. To find the location of this zero-twist slice for each subject, a linear gradient from base to apex was assumed for the twist, resulting in a linear interpolation equation of twist as a function of distance from the base. This enabled the calculation of the subject-specific location of the zero-twist slice along the z-axis. By knowing the location of the zero-twist slice, **Eq. 3a** can be used to find the absolute shear angle, where *φ* is substituted by zero for the zero-twist slice and *D* is the distance of each short-axis slice from the zero-twist slice. Once *θ*_CL_ was obtained, **Eq. 3b** was re-arranged to solve for E_CL_ for each segment. This resulted in an AHA 17 segment model for E_CL_ for each subject.

Finally for the third and fourth steps, the remaining two shear components of the Green strain E_CR_ and E_LR_ were calculated from the polar strain and principal strains of the short-axis and long-axis images, respectively. Furthermore, from 2D continuum mechanics, the off-diagonal shear strain in each 2D plane (i.e., short-axis and long-axis images) is related to its respective polar and principal strain via the following two equations:

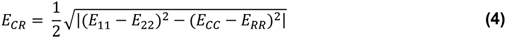

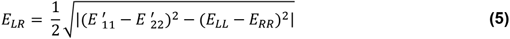

To avoid negative radicands, the absolute value of the difference was taken as shown in **Eqs. 4** and **5**. Completion of steps 3 and 4 resulted in E_CR_ and E_LR_ for each segment in the AHA 17-segment model.

### 3D Strain Attributes

**Fig. 6.**
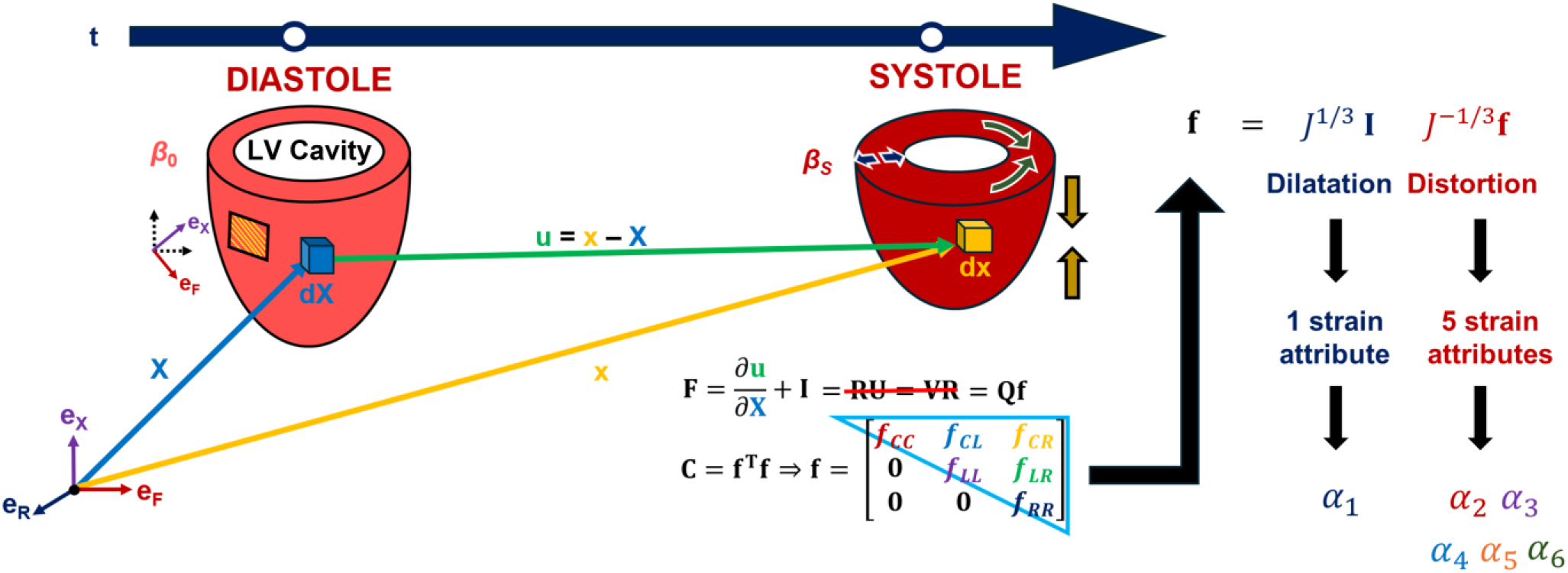
The theoretical workflow to obtain the strain attributes. From the voxel displacement field, the local deformation gradient tensor **F** is obtained. From **F**, the right Cauchy-Green stretch tensor **C** is obtained, which is used to obtain the upper triangular stretch tensor **f**. The separation of **f** into dilatation and distortion allows for the calculation of all six strain attributes.

From the previous section, each component of the full 3D Green strain tensor in **Eq. 1** was obtained for each segment in the AHA 17-segment model (i.e., E_CC_, E_LL_, E_RR_, E_CL_, E_CR_, and E_LR_) over the cardiac cycle. To find the strain attributes, the following three steps were conducted: (1) finding the material axes over the region of interest (ROI); (2) calculating the upper triangular stretch tensor **f** from the 3D Greens strain tensor via **Eq. 6** [41]; and (3) calculating each strain attribute from their respective upper triangular stretch via **Eq. 7**. For step 1, the ROI was defined over the full thickness of the myocardium; hence, the mean material axes aligned with the polar axes as the mean fiber direction was assumed to be circumferentially aligned over the full myocardial thickness. This implied that the mean fiber direction aligned with the circumferential direction; the in-plane perpendicular or the cross-fiber direction aligned with the longitudinal direction; and out-of-plane perpendicular or the material radial direction aligned with the polar radial direction (shown in **Figure 6**). For step 2, the components of the upper triangular stretch tensor can be calculated directly from the component of the Green strain tensor using **Eq. 6** as the mean material directions align with the polar axes [41].

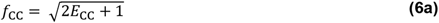

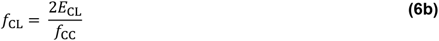

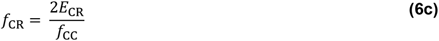

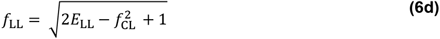

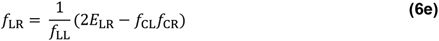

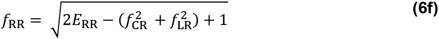

For step 3, strain attributes were calculated directly from the components of **f**. However, it is important to note that the deformation captured by the **f** was separated into a dilatation (i.e., deformation that is *isomorphic* or a change in size) and a distortion (i.e., deformation that is *isochoric* or a change in shape) using Flory’s decomposition [59]. Separating the deformation this way would delineate the changes in the myocardial volume from the changes in the shape of the myocardial mesostructure. To achieve this, **f** is decomposed into a dilatational part (*J*^1/3^ **I**) and a distortional part (*J* ^-1/3^ **f**), respectively, via **f** = (*J*^1/3^ **I**) (*J* ^-1/3^ **f**), where *J* is the Jacobian or the incompressibility constraint, defined as *J* = det(**F**) = det(**f**) = *f*_CC_ *f*_LL_ *f*_RR_. From there, the strain attributes can be defined from the components of both tensors as shown below in **Eq. 7** [24,41].

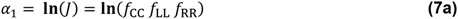

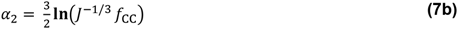

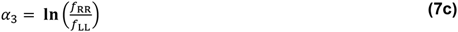

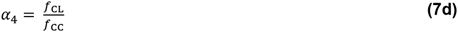

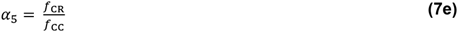

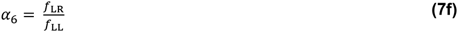

where *α*_1_ is the dilatational strain attribute, *α*_2_ is the mean fiber strain attribute, *α*_3_ is the mean laminar thickening strain attribute, *α*_4_ is the shear strain attribute due to mean laminar sliding, *α*_5_ is the shear strain attribute due to mean laminar tilting about the mean cross-fiber axis, and *α*_6_ is the shear strain attribute due to mean laminar tilting about the mean fiber axis [24,41] . To achieve mean fiber strains consistent with what was reported in previous studies that measured fiber strains, the 3/2 factor in **Eq. 7b** was dropped. From **Eq. 7**, *α*_1_ is the strain attribute that quantifies myocardial dilatation while *α*_2_, *α*_3_, *α*_4_, *α*_5_, and *α*_6_ quantify myocardial distortion.

Because these strain attributes were derived using the 3D Green strain tensor obtained from the previous section, each strain attribute was obtained for each segment of the AHA 17-segment model that time-varies over the cardiac cycle.

### Isochoric Green Strain

Because the strain attributes were separated into dilatation and distortion, the Green strain was, likewise, separated to observe if this separation enhances the regional description of the Green strain relative to the myocardial architecture. Also, the isochoric part of the Green Strain is commonly used in biomechanical modeling of the myocardium. Past studies of MRI-based estimations of the Green strain did not routinely separate the Green strain as such an estimation would require the measurement of *J*, which is not straightforward from **F** [34–36,44,52,60]. However, because this work employs the upper triangular stretch tensor, its determinant is relatively straightforward as shown in the previous section. Hence, this enabled the calculation of the isochoric Green strain. Since the isochoric part of **F** is 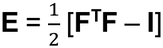, the isochoric part of **E** is 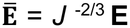, where 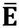 is the isochoric Green strain. Using this equation, the isochoric Green strain was calculated using the previously obtained 3D Green strain tensor, resulting in an AHA 17-segment model for each component of the isochoric Green strain per each subject that time-varies with the cardiac cycle.

### Engineering Strain

To calculate the 3D engineering strain, the definition of the Biot strain was used, where **ε** = **U** – **I**. **U** is the right Cauchy-Green stretch tensor (calculated from **E** via 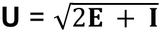), **I** is the identity tensor, and **ε** is the Biot strain. This is because the Biot strain is easier to calculate than the engineering strain due to the latter’s dependence on **F** (which is not possible to calculate from **E** without assuming additional constraints), and the Biot strain and the engineering strain are equal when there are no rigid rotations. This condition is satisfied in MRI acquisition as the acquired imaging slices do not rigidly rotate in space. Also, in 1D, the Biot strain is equal to (*L* – *L*_0_)/*L*_0_ = Δ*L*/*L_0_*, which achieves the same definition sought after in this section — that is, the definition used clinically and implemented by several commercial strain analysis packages. This yields segmental strains that vary over the cardiac cycle for each segment in of the AHA 17-segment model for each subject.

### Strain Validation

To validate the Green strains, we compared our values to values obtained in a multicenter reproducibility study [36]. The Green strains obtained in this study were within the standard deviations reported in the multicenter study, where the reported strains in the study were E_CC_ = −0.18 ± 0.03, E_LL_ = −0.15 ± 0.02, and E_RR_ = 0.35 ± 0.16 [36]. To validate the strain attributes, the relationship between the Green strains and their respective derived strain attributes obtained herein from 2D DENSE was compared with the Green strains and respective strain attributes obtained from the 3D DENSE strains reported in Figure 4 of Zhong et al.’s paper [35]. Zhong et al.’s strains were used since all six Green strain components are needed to calculate the strain attributes, and it is one of the few DENSE studies that does so. Furthermore, the 3D DENSE mid-myocardial Green strains reported in Zhong et al. were reproduced herein, and their respective mid-myocardial strain attributes were derived using **Eq. 6** and **Eq. 7** [35]. Likewise, the mid-myocardial Green strains and respective strain attributes were calculated from the 2D DENSE dataset acquired in this work (as already demonstrated in a previous section).

To quantify the relationship between the Green strains and their respective strain attributes, the ratio between each Green strain component and its corresponding strain attribute counterpart at end-systole was calculated (e.g., E_CC_ / *α*_2_, E_RR_ / *α*_3_, and E_LR_ / *α*_6_). This was done for both the 3D DENSE strains from Zhong et al. and the 2D DENSE strains in the work herein [35]. Finally, to compare the similarity in the ratios between both studies, two steps were needed. First, the percent deviation between both ratios were quantified via **Eq. 8** below

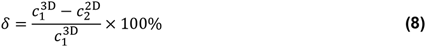

where 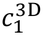 is ratio between a Green strain component and its respective strain attribute from 3D DENSE (e.g., E_CC_ / *α*_2_), 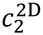 is the scaling coefficient between a Green strain component and its respective strain attribute from 2D DENSE, and *ẟ* is the percent deviation of 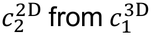. Second, the percent similarity between both ratios was quantified using *ρ* = 1 − *ẟ*, where *ρ* is the percent similarity between 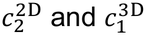.

### Statistics

For each subject, the aforementioned methodology in the previous 4 sections resulted in time-varying segmental strains for each component of the Green strain tensor, the isochoric Green strain tensor, engineering strain, and strain attributes. To enable mean calculations over the 10 subjects, the number of frames were interpolated to 57 frames to ensure a consistent temporal resolution across all subjects. For every strain tensor component per strain definition, the mean global strain, mean regional strain, and mean peak systolic strains were calculated. To calculate the global strain, two steps were done for each strain component: (1) each segmental strain was averaged over all 10 subjects; and (2) the resulting mean segmental strains were averaged over all 16 segments of the AHA bullseye. This produced a global temporal curve for each strain tensor component representing its variation over the cardiac cycle.

To calculate the mean regional strain per strain component (i.e., for the base, mid, and apical slices), the basal segmental strains were averaged to produce the mean basal strain per subject, which was then averaged over all 10 subjects to produce the mean basal strain. Likewise, this process was done for the mid and apical segmental strains as well. Finally, for the peak systolic strains, the strain values at the end-systolic frames for each component of the Green strain and strain attributes were used to produce the AHA bullseye plots for each strain component per subject. The resulting peak systolic AHA bullseye plots were averaged over all 10 subjects.

For each mean that was calculated over all 10 subjects, the standard deviation was calculated as well. For the global strains, to test the hypothesis of whether the difference between the means was significant, a one-way ANOVA was used to assess whether the difference between all 4 global strains (Green strain, strain attributes, isochoric Green strain, and the engineering strain) was significant. This means that for any given strain component, 4 global temporal curves (one curve per strain definition) were used for the one-way ANOVA. For the regional strain, a paired *t*-test was used to assess whether the difference between each component of the Green and its analogous strain attribute was significant in each region (i.e., the base, mid, and apex). For the AHA bullseye, a paired *t*-test was used for each segment in the bullseye to assess whether the difference in the segmental peak systolic strains between each component of the Green strain and its analogous strain attribute was significant. For all of these statistical tests, the threshold of significance for the *p*-value was 0.05. Wherever multiple comparisons were needed, a Bonferroni correction was used to correct the threshold of significance in accordance with the number of pair-wise comparisons required.

## RESULTS

DENSE data was acquired successfully in all 10 subjects. However, due to the presence of artifacts in the basal slice in 3 subjects, their basal *α*_1_, *α*_4_, and *E*_CL_ were excluded from the analysis. The results section is organized into 4 main subsections: (1) global strain results, (2) regional strain results, (3) AHA 17-segment strain results, and (4) myocardial compressibility results. The global strains show how both the normal and shear strains varied with time, where all 4 strain definitions introduced herein (Green strain, strain attributes, isochoric Green strains, and engineering strain) were compared. The regional strain results show the base-to-apex transition of both the Green strains and strain attributes. The AHA 17-segment strain results elucidate how each component of the Green strain and strain attributes varied spatially within the myocardium and how the respective segments compare between both strain definitions. Finally, the myocardial compressibility results show the temporal variation, base-to-apex transition, and AHA 17-segment bullseye of both *J* and *α*_1_. These results help answer the central question of this study: *How do the polar geometric strains compare with mesostructure-based strains in quantifying myocardial deformation?*

### Global Strains

#### Normal Strains

As stated before, the normal strains represent the diagonal components of the strain tensor. Hence, the global values for the diagonal components of the myocardial Green strain (*E*_CC_, *E*_RR_, and *E*_LL_), strain attributes (*α*_2_ and *α*_3_), isochoric Green strains (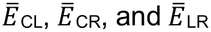), and the engineering strains (*ε*_CC_, *ε*_RR_, and *ε*_LL_) along their respective axial directions are shown below in **Figure 7**, with the associated numerical values shown in **Table 1**. Because *α*_3_ is a ratio of radial thickening to cross-fiber shortening, it was not displayed in the longitudinal plot in **Figure 7** due to being positive, compared to the negative longitudinal strains. Along the circumferential direction (which is also the mean fiber direction), *α*_2_ predicted a significantly lower magnitude of deformation compared to *E*_CC_ (*p* < 0.01), E̅_CC_ (*p* < 0.001), and *ε*_CC_ (*p* < 0.001). On the contrary, along the radial direction, *α*_3_ predicted a significantly higher magnitude of deformation compared to *E*_RR_ (*p* < 0.001) and *ε*_RR_ (*p* < 0.001) but a comparable magnitude relative to E̅_RR_. *This implies that the strains based on mesostructural directions predict deformation patterns significantly different to strains based on polar geometrical directions*.

**Fig. 7.**
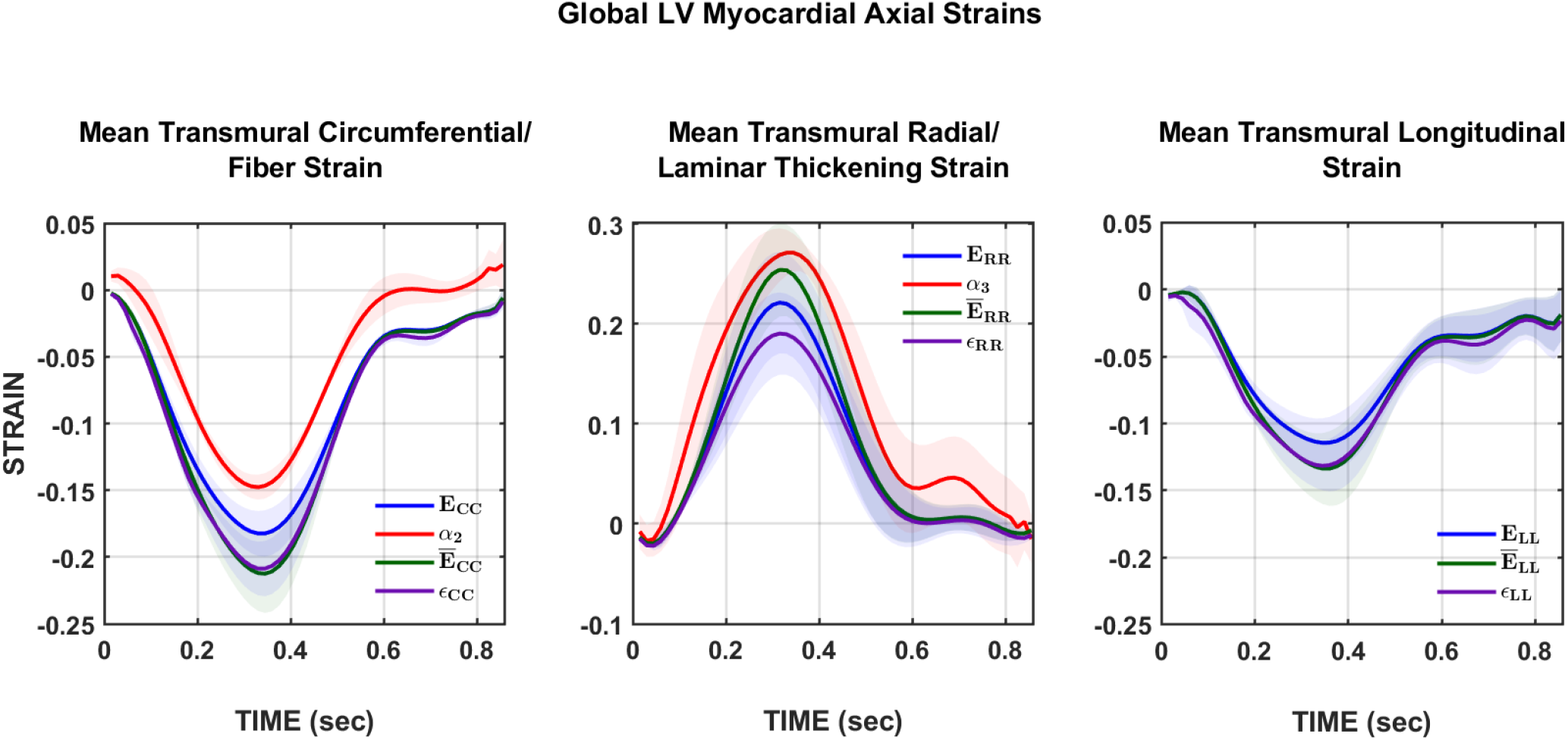
Temporal evolution of normal global strains in the LV myocardium across a normalized cardiac cycle. The panels show the strains along the circumferential/mean fiber direction (left), radial direction/laminar thickening (middle), and longitudinal direction (right). The curves displayed are the Green strains (blue), strain attributes (red), isochoric Green strains (green), and the engineering strains (purple).

**Table 1.** End-systolic basal, mid, apical, and global strains (mean ± SD) for the Green strain (**E**), strain attributes (*α*), isochoric Green strain (E̅), and engineering strain (**ε**).

| **Green Strain (E)** |  |  |  |  |  |  |
|---|---|---|---|---|---|---|
| **Base** | *E*<sub>CC</sub> | (−17 ± 1.8) % | *E*<sub>RR</sub> | (28 ± 8.0) % | *E*<sub>LL</sub> | (−9.4 ± 2.0) % |
|  | *E*<sub>CL</sub> | (−4.3 ± 0.9) % | *E*<sub>CR</sub> | (7.4 ± 1.9) % | *E*<sub>LR</sub> | (14 ± 2.4) % |
| **Mid** | *E*<sub>CC</sub> | (−18 ± 2.9) % | *E*<sub>RR</sub> | (19 ± 4.9) % | *E*<sub>LL</sub> | (−13 ± 2.9) % |
|  | *E*<sub>CL</sub> | (4.1 ± 2.0) % | *E*<sub>CR</sub> | (6.6 ± 1.0) % | *E*<sub>LR</sub> | (9.4 ± 2.2) % |
| **Apex** | *E*<sub>CC</sub> | (−20 ± 2.3) % | *E*<sub>RR</sub> | (19 ± 11) % | *E*<sub>LL</sub> | (−12 ± 2.9) % |
|  | *E*<sub>CL</sub> | (2.4 ± 0.6) % | *E*<sub>CR</sub> | (7.0 ± 1.2) % | *E*<sub>LR</sub> | (12 ± 1.6) % |
| **Global** | *E*<sub>CC</sub> | (−18 ± 1.7) % | *E*<sub>RR</sub> | (22 ± 5.1) % | *E*<sub>LL</sub> | (−11 ± 1.7) % |
|  | *E*<sub>CL</sub> | (1.3 ± 3.7) % | *E*<sub>CR</sub> | (7.0 ± 0.4) % | *E*<sub>LR</sub> | (12 ± 2.2) % |
| **Strain Attributes (*α<sub>i</sub>*)** |  |  |  |  |  |  |
| **Base** | *α*<sub>1</sub> | (−17 ± 7.9) % | *α*<sub>2</sub> | (−15 ± 2.6) % | *α*<sub>3</sub> | (28 ± 7.1) % |
|  | *α*<sub>4</sub> | (−13 ± 2.5) % | *α*<sub>5</sub> | (22 ± 5.1) % | *α*<sub>6</sub> | (37 ± 6.1) % |
| **Mid** | *α*<sub>1</sub> | (−26 ± 9.8) % | *α*<sub>2</sub> | (−14 ± 1.8) % | *α*<sub>3</sub> | (29 ± 5.8) % |
|  | *α*<sub>4</sub> | (13 ± 7.3) % | *α*<sub>5</sub> | (21 ± 4.6) % | *α*<sub>6</sub> | (23 ± 5.6) % |
| **Apex** | *α*<sub>1</sub> | (−31 ± 11) % | *α*<sub>2</sub> | (−16 ± 4.8) % | *α*<sub>3</sub> | (24 ± 9.9) % |
|  | *α*<sub>4</sub> | (8.2 ± 2.2) % | *α*<sub>5</sub> | (23 ± 5.3) % | *α*<sub>6</sub> | (30 ± 3.0) % |
| **Global** | *α*<sub>1</sub> | (−25 ± 6.9) % | *α*<sub>2</sub> | (−15 ± 0.9) % | *α*<sub>3</sub> | (27 ± 2.3) % |
|  | *α*<sub>4</sub> | (4.6 ± 12) % | *α*<sub>5</sub> | (22 ± 1.2) % | *α*<sub>6</sub> | (30 ± 7.0) % |
| **Isochoric Green Strain (Ē)** |  |  |  |  |  |  |
| **Base** | *Ē*<sub>CC</sub> | (−19 ± 2.0) % | *Ē*<sub>RR</sub> | (31 ± 8.8) % | *Ē*<sub>LL</sub> | (−10 ± 2.2) % |
|  | *Ē*<sub>CL</sub> | (−4.7 ± 1.0) % | *Ē*<sub>CR</sub> | (8.2 ± 2.1) % | *Ē*<sub>LR</sub> | (15 ± 2.7) % |
| **Mid** | *Ē*<sub>CC</sub> | (−21 ± 3.5) % | *Ē*<sub>RR</sub> | (23 ± 5.8) % | *Ē*<sub>LL</sub> | (−15 ± 3.3) % |
|  | *Ē*<sub>CL</sub> | (−4.9 ± 2.4) % | *Ē*<sub>CR</sub> | (7.8 ± 1.2) % | *Ē*<sub>LR</sub> | (11 ± 2.6) % |
| **Apex** | *Ē*<sub>CC</sub> | (−24 ± 2.9) % | *Ē*<sub>RR</sub> | (23 ± 14) % | *Ē*<sub>LL</sub> | (−15 ± 3.3) % |
|  | *Ē*<sub>CL</sub> | (2.9 ± 0.7) % | *Ē*<sub>CR</sub> | (8.4 ± 1.5) % | *Ē*<sub>LR</sub> | (15 ± 2.0) % |
| **Global** | *Ē*<sub>CC</sub> | (−21 ± 2.8) % | *Ē*<sub>RR</sub> | (25 ± 4.6) % | *Ē*<sub>LL</sub> | (−13 ± 2.5) % |
|  | *Ē*<sub>CL</sub> | (1.0 ± 5.0) % | *Ē*<sub>CR</sub> | (8.1 ± 0.3) % | *Ē*<sub>LR</sub> | (14 ± 2.3) % |
| **Engineering Strain (ε)** |  |  |  |  |  |  |
| **Base** | ε<sub>CC</sub> | (−19 ± 2.2) % | ε<sub>RR</sub> | (24 ± 6.8) % | ε<sub>LL</sub> | (−11 ± 1.9) % |
|  | ε<sub>CL</sub> | (−5.7 ± 1.1) % | ε<sub>CR</sub> | (7.7 ± 1.9) % | ε<sub>LR</sub> | (13 ± 2.5) % |
| **Mid** | ε<sub>CC</sub> | (−20 ± 3.8) % | ε<sub>RR</sub> | (17 ± 4.2) % | ε<sub>LL</sub> | (−14 ± 3.3) % |
|  | ε<sub>CL</sub> | (4.7 ± 2.5) % | ε<sub>CR</sub> | (6.5 ± 1.2) % | ε<sub>LR</sub> | (9.1 ± 2.2) % |
| **Apex** | ε<sub>CC</sub> | (−23 ± 3.2) % | ε<sub>RR</sub> | (16 ± 10) % | ε<sub>LL</sub> | (−14 ± 3.2) % |
|  | ε<sub>CL</sub> | (2.4 ± 0.8) % | ε<sub>CR</sub> | (7.1 ± 1.4) % | ε<sub>LR</sub> | (12 ± 1.4) % |
| **Global** | ε<sub>CC</sub> | (−21 ± 2.0) % | ε<sub>RR</sub> | (19 ± 4.0) % | ε<sub>LL</sub> | (−13 ± 1.7) % |
|  | ε<sub>CL</sub> | (1.1 ± 4.5) % | ε<sub>CR</sub> | (7.1 ± 0.6) % | ε<sub>LR</sub> | (11 ± 2.1) % |
| **The Jacobian (*J*)** |  |  |  |  |  |  |
|  | ***Base*** | 0.86 ± 0.060 | ***Mid*** | 0.79 ± 0.074 | ***Apex*** | 0.76 ± 0.060 |
|  |  |  | ***Global*** | 0.80 ± 0.054 |  |  |

The magnitude of each strain definition at end-systole is displayed in **Figure 7**, and the respective numerical values can be found in **Table 1**. By comparing the relative magnitudes of the strain attributes, the global end-systolic *α*_3_ was roughly double the magnitude of the global end-systolic *α*_2_. However, for the Green strain, the global end-systolic E_RR_ was roughly comparable in magnitude with the global end-systolic E_CC_. Likewise, the relative magnitudes of the isochoric Green strains for 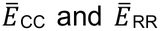 were roughly comparable as well. Finally, for the engineering strains, *ε*_CC_ predicts a significantly higher deformation compared to either *E*_CC_ (*p* < 0.001) or *α*_2_ (*p* < 0.001) whereas *ε*_RR_ predicts a significantly lower deformation compared to either *E*_RR_ (*p* < 0.001) or *α*_3_ (*p* < 0.001). Similarly, *ε*_LL_ predicts a significantly higher deformation compared to *E*_LL_ (*p* < 0.001). This means that engineering strain, which is defined for infinitesimal deformations, predicts significantly different strain patterns from those defined for finite deformations.

**Fig. 8.**
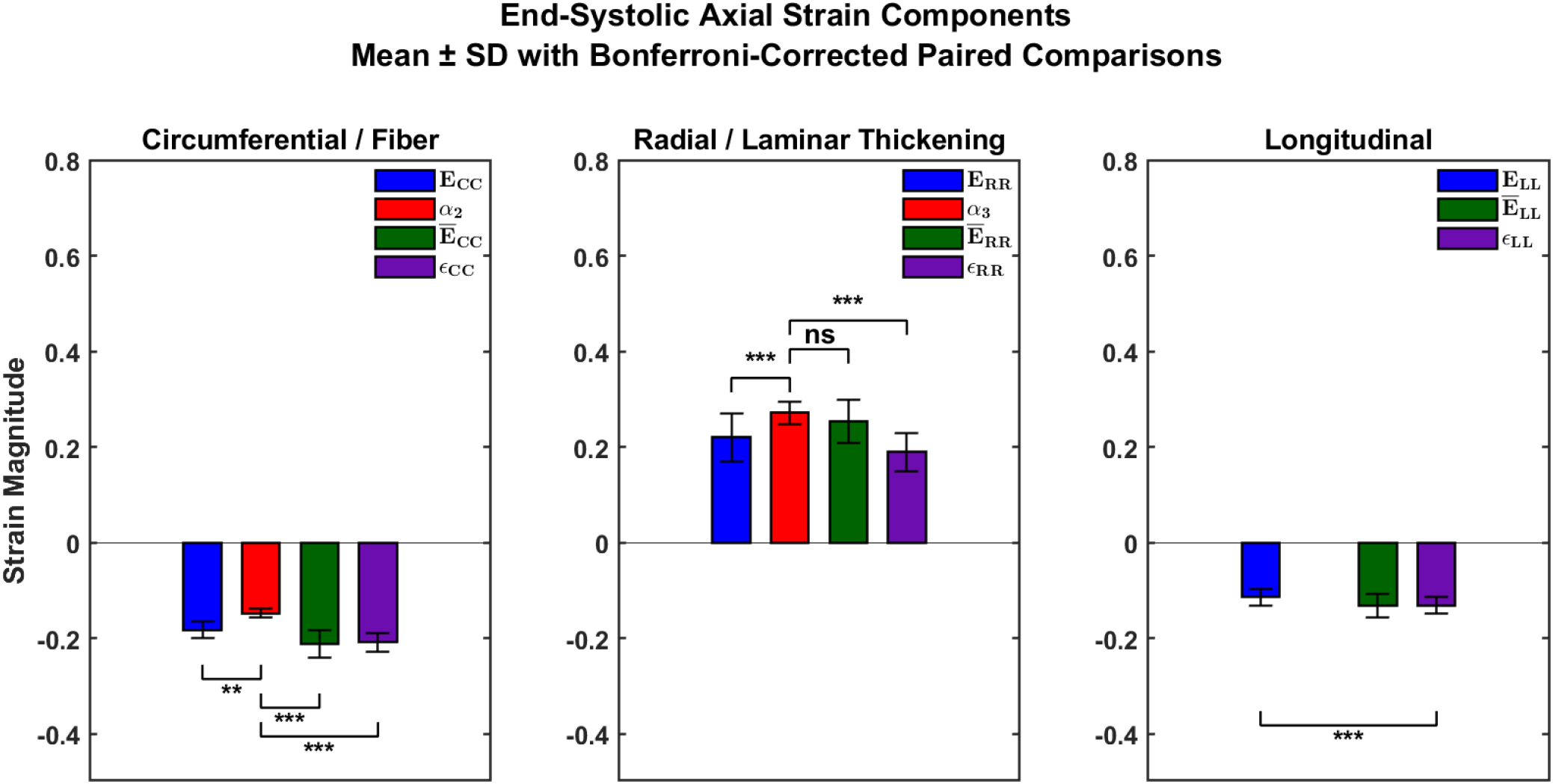
End-systolic Normal Strains (Mean ± SD) for the Green strain (**E**), strain attributes (*α*i), isochoric Green strain (E̅), and engineering strain (**ε**). * indicates *p* < 0.05, ** indicates *p* < 0.01, *** indicates *p* < 0.001, while ns indicates not significant.

#### Shear Strains

As stated before, the shear strains represent the off-diagonal components of the strain tensor. Hence, the global values for the shear components of the myocardial Green strain (*E*_CL_, *E*_CR_, and *E*_LR_), strain attributes (*α*_4_, *α*_5_, and *α*_6_), isochoric Green strain (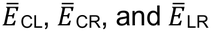), and the engineering strain (*ε*_CL_, *ε*_CR_, and *ε*_LR_) are shown below in **Figure 9**, and the respective numerical values can be found in **Table 1**. From the graphs in **Figure 9**, it is clear that the shear deformation predicted by the strain attributes is greater than that of the other 3 strain definitions across all 3 shear modes. Furthermore, for the shear mode of laminar twist (i.e., shearing of the circumferential-longitudinal plane), *α*_4_ predicted a comparable magnitude of deformation relative to *E*_CL_ and 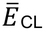 but a significantly higher magnitude of deformation compared to *ε*_CL_ (*p* < 0.05). For the shear mode of laminar imbrication due to tilting about the longitudinal axis (i.e., shearing of the circumferential-radial plane), *α*_5_ predicted a significantly higher magnitude of deformation compared to *E*_CR_ (*p* < 0.001), 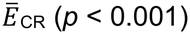, and *ε*_CR_ (*p* < 0.001). Finally, for the shear mode of laminar imbrication due to tilting about the circumferential axis (i.e., shearing of the longitudinal-radial plane), *α*_6_ predicted a significantly higher magnitude of deformation compared to *E*_LR_ (*p* < 0.001), 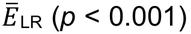, and *ε*_LR_ (*p* < 0.001). Once again, this means that the deformation ‘seen’ from the mesostructural directions is significantly different to that seen from the polar geometrical directions.

**Fig. 9.**
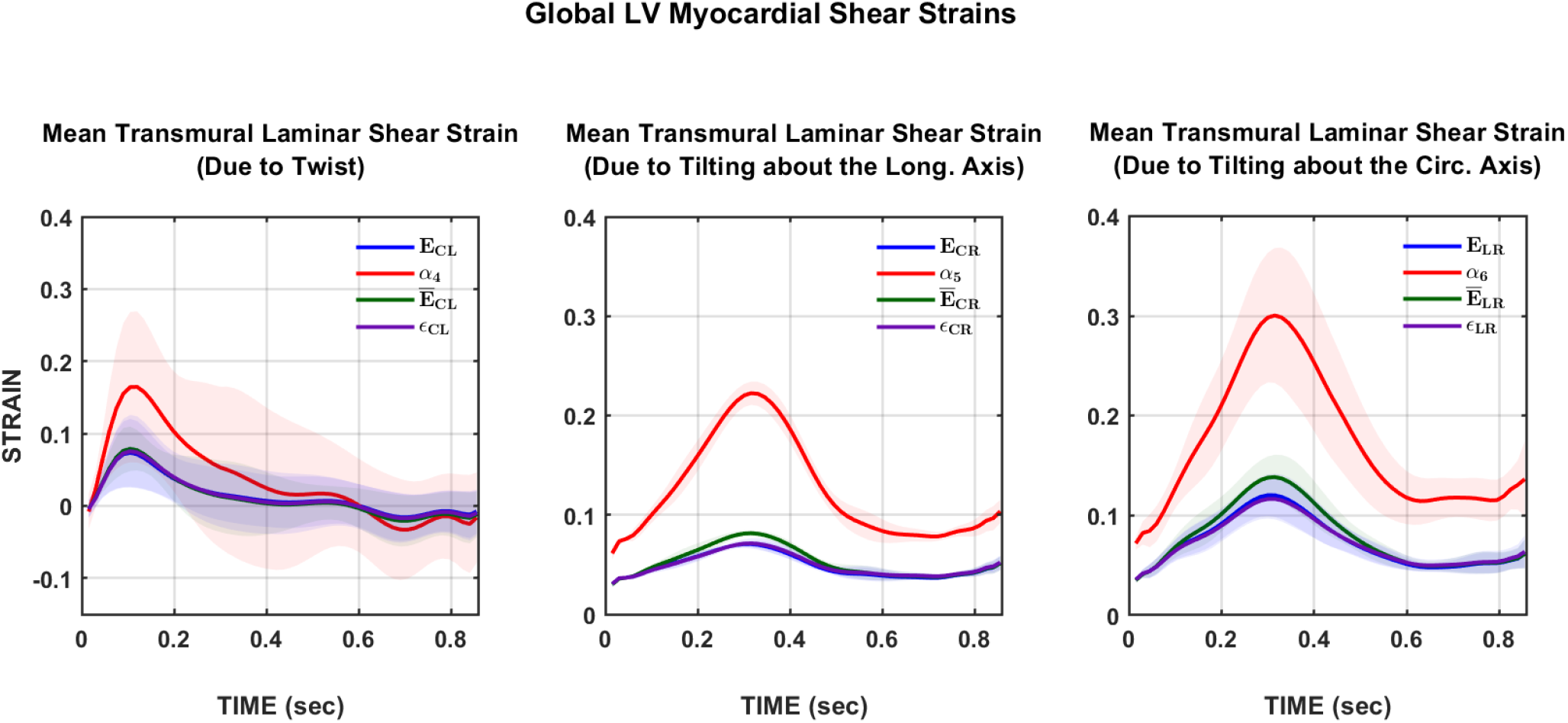
Temporal evolution of shear global strains in the LV myocardium across a normalized cardiac cycle. The panels show the strains pertaining to the circumferential–longitudinal shear (left), circumferential–radial shear (middle), and longitudinal–radial shear (right). The curves displayed are the Green strains (blue), strain attributes (red), isochoric Green strains (green), and the engineering strains (purple).

Regarding the relative magnitudes of the 3 shear modes within each strain definition, the end-systolic shear strains are displayed in **Figure 10**, and the respective numerical values can be found in **Table 1**. By comparison, the relationship between the 3 shear modes is the same across all 4 strain definitions: the longitudinal–radial shear mode has the greatest magnitude, followed by the circumferential–radial shear mode, which is followed by the circumferential– longitudinal shear and has the lowest magnitude. Hence, ordering the shear terms for each strain definition from lowest to highest results in *E*_CL_ < *E*_CR_ < *E*_LR_ for the Green strain (*p* < 0.001), *α*_4_ < *α*_5_ < *α*_6_ for the strain attributes, E̅_CL_ < E̅_CR_ < E̅_LR_ for the isochoric Green strain (*p* < 0.001), and *ε*_CL_ < *ε*_CR_ < *ε*_LR_ for the engineering strain (*p* < 0.001). For engineering strain, while *ε*_CR_ was comparable to *E*_CR_, *ε*_CL_ was significantly different from *E*_CL_ (*p* < 0.001), and *ε*_LR_ was significantly different from *E*_LR_ (*p* < 0.01).

**Fig. 10.**
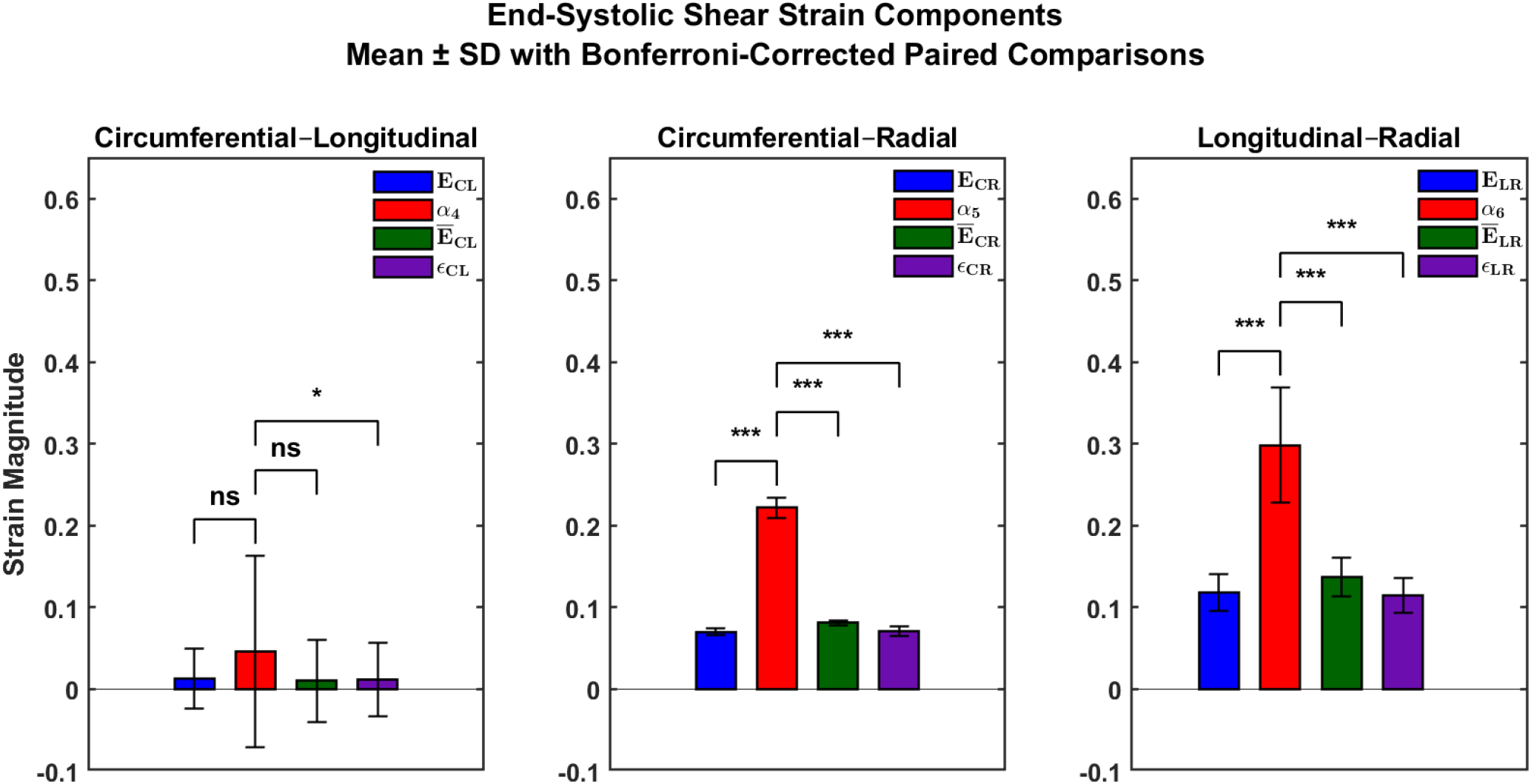
End-systolic Shear Strains (Mean ± SD) for the Green strain (**E**), strain attributes (*α*), isochoric Green strain (E̅), and engineering strain (**ε**).

### Global Regional Strains (Base to Apex)

Given the similarity of the isochoric Green strain in capturing the mesostructural deformation of the myocardium to the Green strain as well as the inapplicability of engineering strain to the myocardium, the focus will be on comparing the Green strain with the strain attributes for the remainder of this section. The Green strain components displayed herein are *E*_CC_, *E*_RR_, and *E*_LR_ and the corresponding strain attributes are *α*_2_, *α*_3_, and *α*_6_. The remaining components are found in the supplemental materials. Hence, the magnitudes of the aforementioned strain components calculated globally at the basal, mid, and apical myocardium are shown in **Figures 11 – 13** in the following subsections. Their respective numerical values are shown in **Table 1**.

#### Circumferential Green Strain (E_CC_) vs. Mean Fiber Strain Attribute (α_2_)

**Figure 11** below shows the base-to-apex transition of both the global *E*_CC_ and *α*_2_. For *E*_CC_, the end-systolic magnitude is comparable between base and mid but is higher at the apex (*p* < 0.001). However, for *α*_2_, the end-systolic magnitude is comparable across the base, mid, and apex. Hence, the increasing order of each strain from base to apex is basal *E*_CC_ ∼ mid *E*_CC_ < apical *E*_CC_ for the circumferential Green strain and basal *α*_2_ ∼ mid *α*_2_ ∼ apical *α*_2_ for the mean fiber strain attribute.

**Fig. 11.**
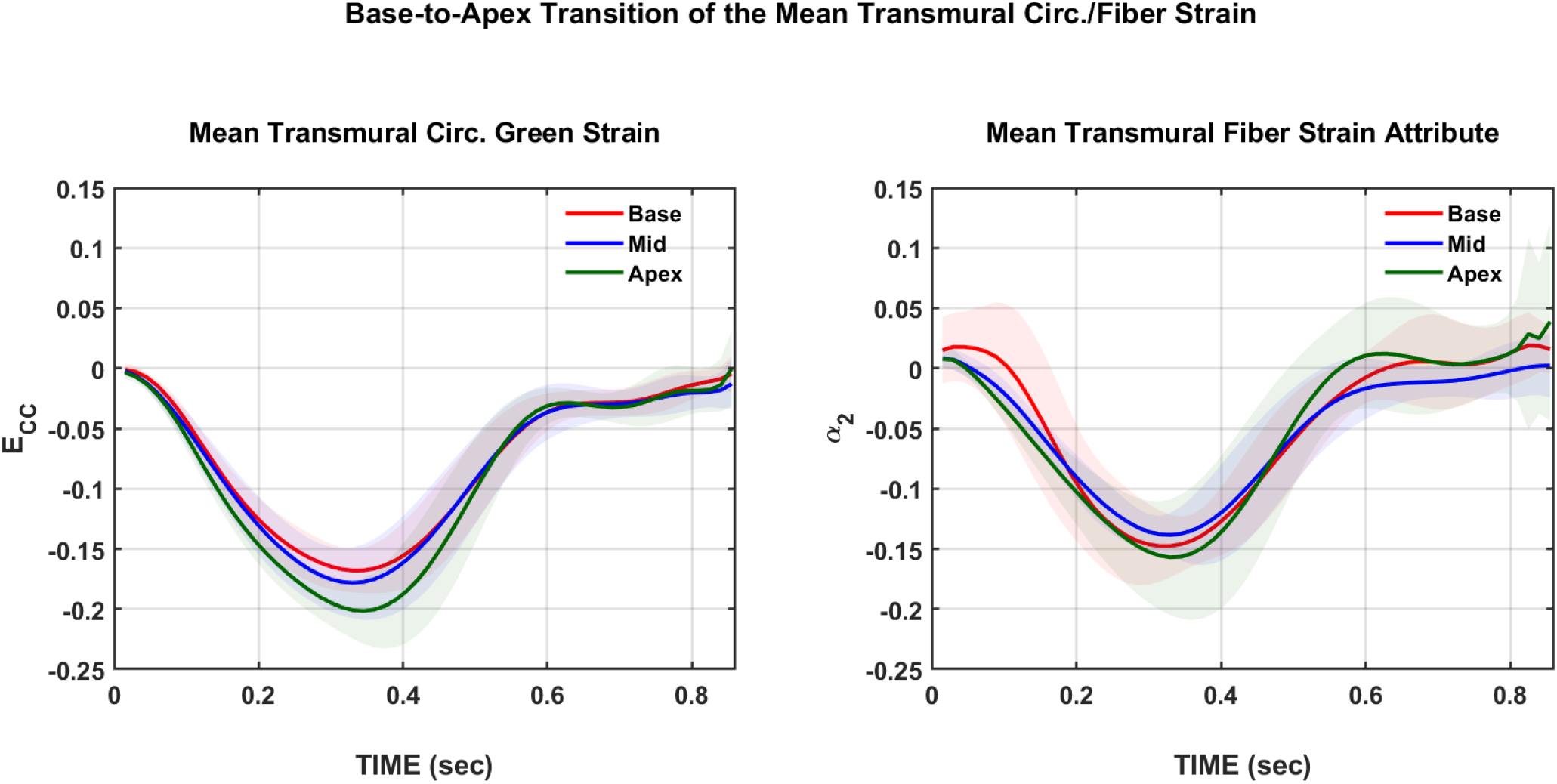
Base-to-apex transition of the strain in the LV myocardium across a normalized cardiac cycle. The panels show the circumferential Green strain (left) and the mean fiber strain attribute (right). The curves displayed pertain to the basal (red), mid (blue), and the apical (green) myocardium.

#### Radial Green Strain (E_RR_) vs. Mean Laminar Thickening Strain Attribute (α_3_)

**Figure 12** below shows the base-to-apex transition of both the global *E*_RR_ and *α*_3_. For *E*_RR_, the end-systolic magnitude is higher at the base (*p* < 0.01) but comparable between the mid and apex. For *α*_3_, the end-systolic magnitude is comparable across the base, mid, and apex. Hence, the increasing order of each strain from base to apex is apical *E*_RR_ ∼ mid *E*_RR_ < basal *E*_RR_ for the circumferential Green strain and apical *α*_3_ ∼ mid *α*_3_ ∼ basal *α*_3_ for the mean laminar thickening strain attribute.

**Fig. 12.**
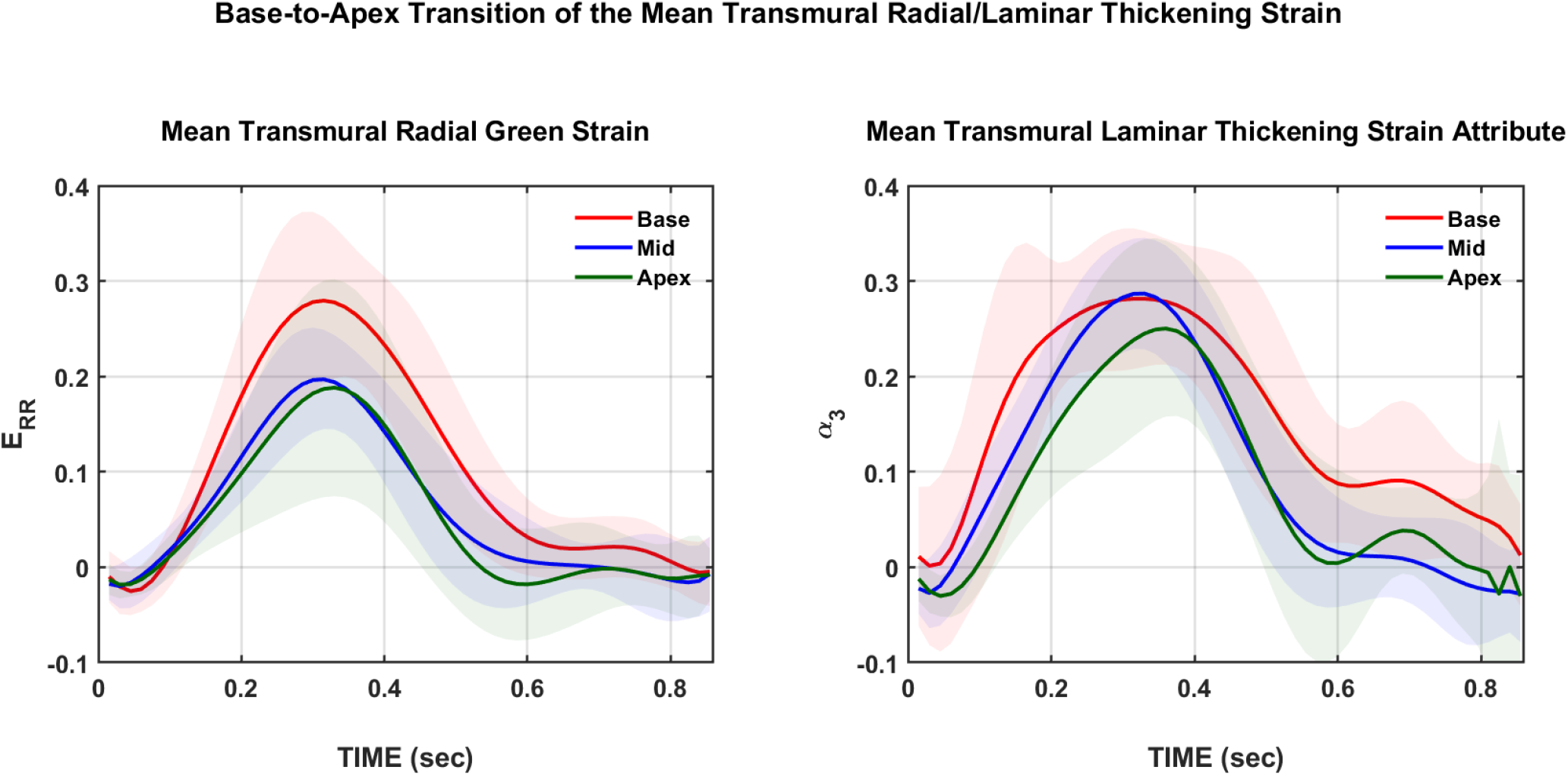
Base-to-apex transition of the strain in the LV myocardium across a normalized cardiac cycle. The panels show the radial Green strain (left) and the mean laminar thickening strain attribute (right). The curves displayed pertain to the basal (red), mid (blue), and the apical (green) myocardium.

#### Longitudinal–Radial Shear Green Strain (E_LR_) vs. Mean Cross-fiber–Radial Strain Attribute (α_6_)

**Figure 13** below shows the base-to-apex transition of both the global *E*_LR_ and *α*_6_. For *E*_LR_, the end-systolic magnitude at the mid is less than that at the base (*p* < 0.0001) and apex (*p* < 0.01), but it is comparable between the base and apex. However, for *α*_6_, while the end-systolic magnitude at the mid is less than that at the base (*p* < 0.0001) and apex (*p* < 0.01), the basal *α*_6_ is greater than the apical *α*_6_ (*p* < 0.01). Hence, the increasing order of each strain from base to apex is mid *E*_LR_ < apical *E*_LR_ ∼ basal *E*_LR_ for the longitudinal–radial shear Green strain and mid *α*_6_ < apical *α*_6_ < basal *α*_6_ for the mean *cross-fiber–radial* strain attribute.

**Fig. 13.**
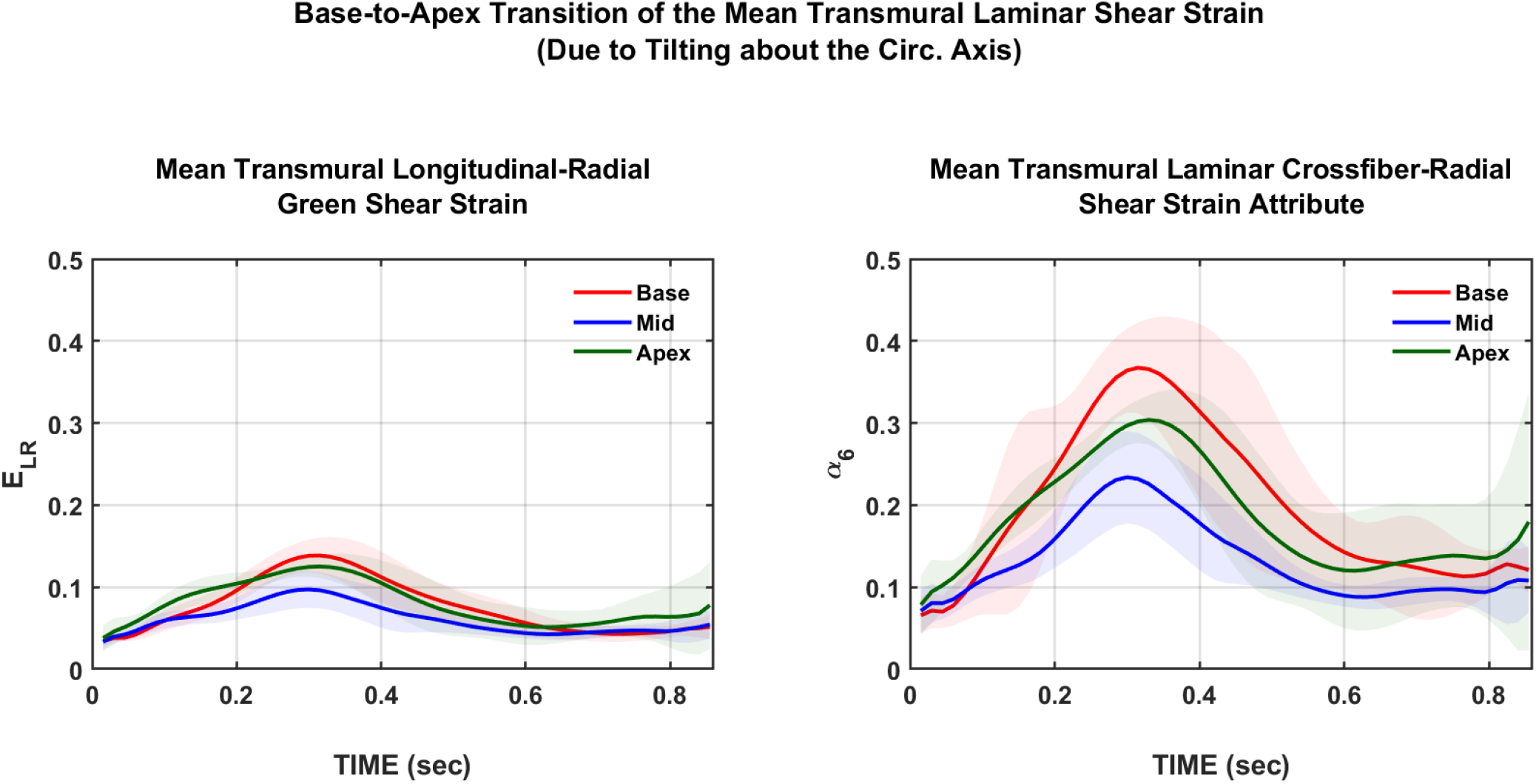
Base-to-apex transition of the strain in the LV myocardium across a normalized cardiac cycle. The panels show the longitudinal–radial shear Green strain (left) and the mean cross-fiber–radial strain attribute (right). The curves displayed pertain to the basal (red), mid (blue), and the apical (green) myocardium.

### AHA 17-Segment Model of the End-systolic Strains

To visualize myocardial strain distribution over the entire LV, the components of both the Green strain (i.e., *E*_CC_, *E*_RR_, and *E*_LR_) and the strain attributes (i.e., *α*_2_, *α*_3_, and *α*_6_) were plotted at end-systole using the AHA 17-segment model, shown in **Figure 14**. Qualitatively, the strain attributes showed greater regional heterogeneity in relation to their respective components of the Green strain, which was a consistent finding across all 10 subjects. For example, the magnitude for the both the *α*_2_ and *α*_3_ were greater in the LV lateral wall than that of the septal wall whereas the difference between both regions is markedly less for both the *E*_CC_ and *E*_RR_. Likewise, *α*_2_ and *α*_3_ were greater in the anterior wall compared to the inferior wall, with a similar distinction for *E*_RR_. For *E*_CC_, the difference between the anterior wall and inferior wall was qualitatively comparable. The regional magnitudes of *α*_6_ were ∼3 – 5 times greater than their analogous regions in the *E*_LR_ bullseye plot. Spatially, *α*_6_ predicted greater heterogeneity such that the LV septal wall was greater in magnitude than the lateral wall. *E*_LR_ predicted far less heterogeneity between both regions.

**Fig. 14.**
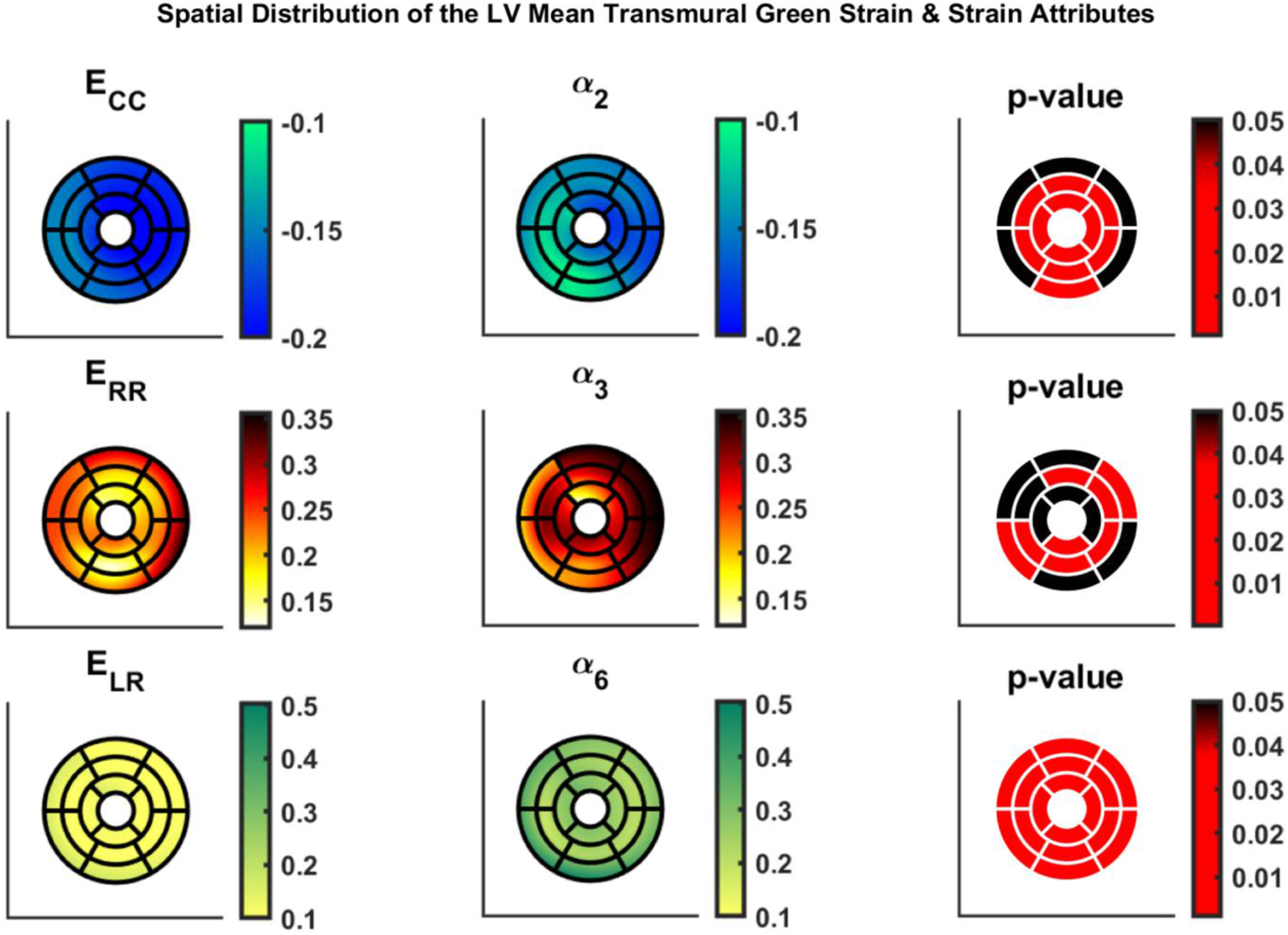
The spatial distribution of select strain components across the AHA 17-segment model in the LV myocardium with the p-values from a segment-to-segment paired t-test shown on the far right panel. Black segments indicate no significance (*p* > 0.05) while red segments indicate differences that were significant (*p* < 0.05).

Quantitatively, the difference between *E*_CC_ and *α*_2_ appeared comparable at the base (except at the basal inferior segment) yet was significant at the mid and apical segments. Similarly, *E*_RR_ and *α*_3_ appeared comparable at the base (with exceptions at the anterolateral and inferoseptal segments) and the apex (except at the inferior segment) but significantly different at the mid (except at the anterolateral segment). However, *E*_LR_ and *α*_6_ appeared significantly different in all the LV segments.

### The Compressibility of the Myocardium

#### The Mean Dilatational Strain & The Jacobian

The global dilatational strain attribute *α* _1_ was plotted over the cardiac cycle in the basal, mid, and apical myocardium (shown in **Figure 15**). A magnitude of *α*_1_ = 0 indicates no change in volume (i.e., incompressible) whereas *α*_1_ ≠ 0 indicates a net change in volume (i.e., compressible) – such that *α*_1_ > 0 indicates volume expansion whereas *α*_1_ < 0 indicates volume shrinkage. Furthermore, the further away *α*_1_ is from the zero line, the greater the amount of compressibility or volume change the myocardium undergoes. From the plot of *α*_1_ in **Figure 15**, the magnitude of the dilatational strain increases from base to mid (*p* < 0.01) but is comparable between the mid and apical regions. To further elucidate this regional relationship, the end-systolic magnitudes *α*1 are shown below in **Figure 16**. Thus, the order of *α*1 in increasing magnitude is basal *α*1 < mid *α*1 ∼ apical *α*1.

In addition, the Jacobian (i.e., measure of compressibility) was plotted over the cardiac cycle in each of the 3 myocardial short-axis slices. Analogous to *α*_1_, a magnitude of *J* = 1 indicates no change in volume (i.e., incompressible) whereas *J* ≠ 1 indicates a net change in volume (i.e., compressible) – such that *J* > 1 indicates volume expansion whereas *J* < 1 indicates volume shrinkage. The further away *J* is from the unity line (*J* = 1), the greater the amount of compressibility or volume change the myocardium undergoes. From the plot of *J* in **Figure 15**, the myocardial compressibility increases from base to mid (*p* < 0.001) but is comparable between the mid and apical regions. Similar to *α*1, the end-systolic magnitudes of *J* are shown below in **Figure 16** (a decreasing magnitude of *J* indicates greater myocardial compressibility and vice versa). Thus, the order of *J* in increasing magnitude is apical *J* ∼ mid *J* < basal *J*. The conclusion drawn from both *J* and *α*_1_ is that (1) the myocardium is *not* incompressible, and (2) myocardial compressibility increases from base to mid but remains relatively comparable between the mid and apex.

**Fig. 15.**
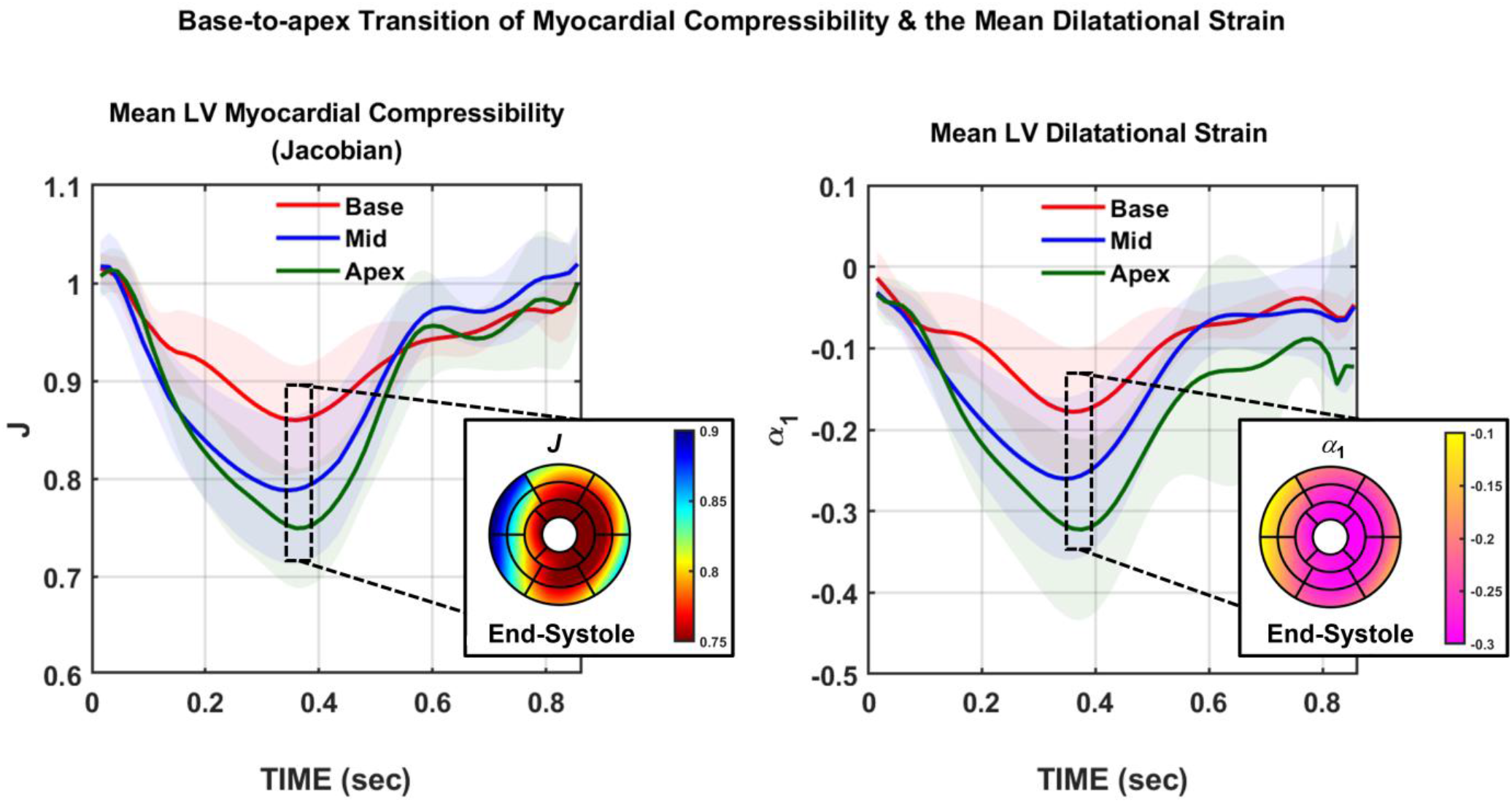
The temporal variation of the Jacobian (left) and α_1_ (right) over the cardiac cycle. Variation is displayed for the basal (red), mid (blue), and apical (green) myocardial short-axis slices. Also shown is the regional distribution of myocardial compressibility (left) and dilatational strain (right) in the AHA 17-segment model at end-systole.

**Fig. 16.**
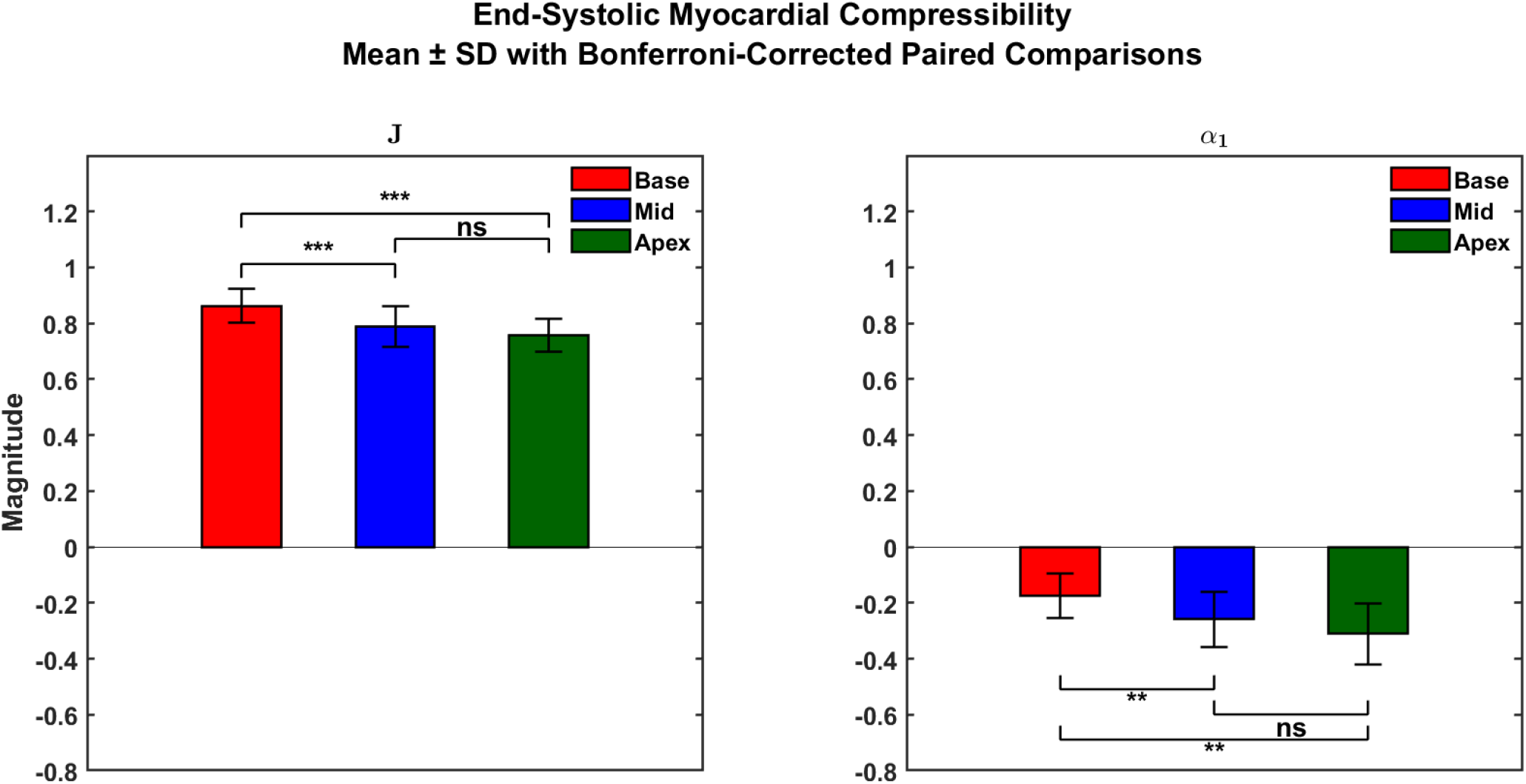
Base-to-apex values (Mean ± SD) for myocardial compressibility (**J**) and dilatational strain attribute (*α*_1_).

Regionally, the spatial distribution of *J* and *α*1 across the LV AHA segments is heterogeneous as shown in bullseye plots in **Figure 15**. Compared to the septal wall, the lateral wall has greater magnitudes of *α*1 as well as both the anterior and inferior walls. Additionally, the relative segmental magnitudes of *α*1 increase from base to apex. A similar pattern can be observed for *J*, where it is greater in the lateral wall than the septal wall, with an increasing base-to-apex gradient. Interestingly, *J* value of 1 did not occur in any segment nor did an *α*1 value of 0, implying no incompressible regions in the myocardium.

### Validation of the Strain Attributes

To validate the strain attributes obtained in this study, we compared our findings to data obtained from Zhong et al. [35]. The first row in **Figure 17** shows the mid-myocardial Green strains reproduced from Figure 4 in Zhong et al. (shown in blue), which were obtained from 3D DENSE. The strain attributes we derived from these 3D DENSE Green strains are shown in red. The second row in **Figure 17** displays the mean mid-myocardial Green strain (shown blue) and derived mean strain attributes (shown in red) from this study, obtained from 2D DENSE. The percent similarity in the scaling coefficients between 2D DENSE and 3D DENSE end-systolic strains was *ρ* = 87.5% for *E*_CC_ and *α*_2_, *ρ* = 96.6% for *E*_RR_ and *α*_3_, and *ρ* = 99.6% for *E*_LR_ and *α*_6_. This indicates that the relationship between the Green strains and their respective strain attributes in the work herein was at least ∼87% similar to the relationship between the Green strains and their respective strain attributes obtained from the 3D DENSE strains reported in Zhong et al. [35].

**Figure 17.**
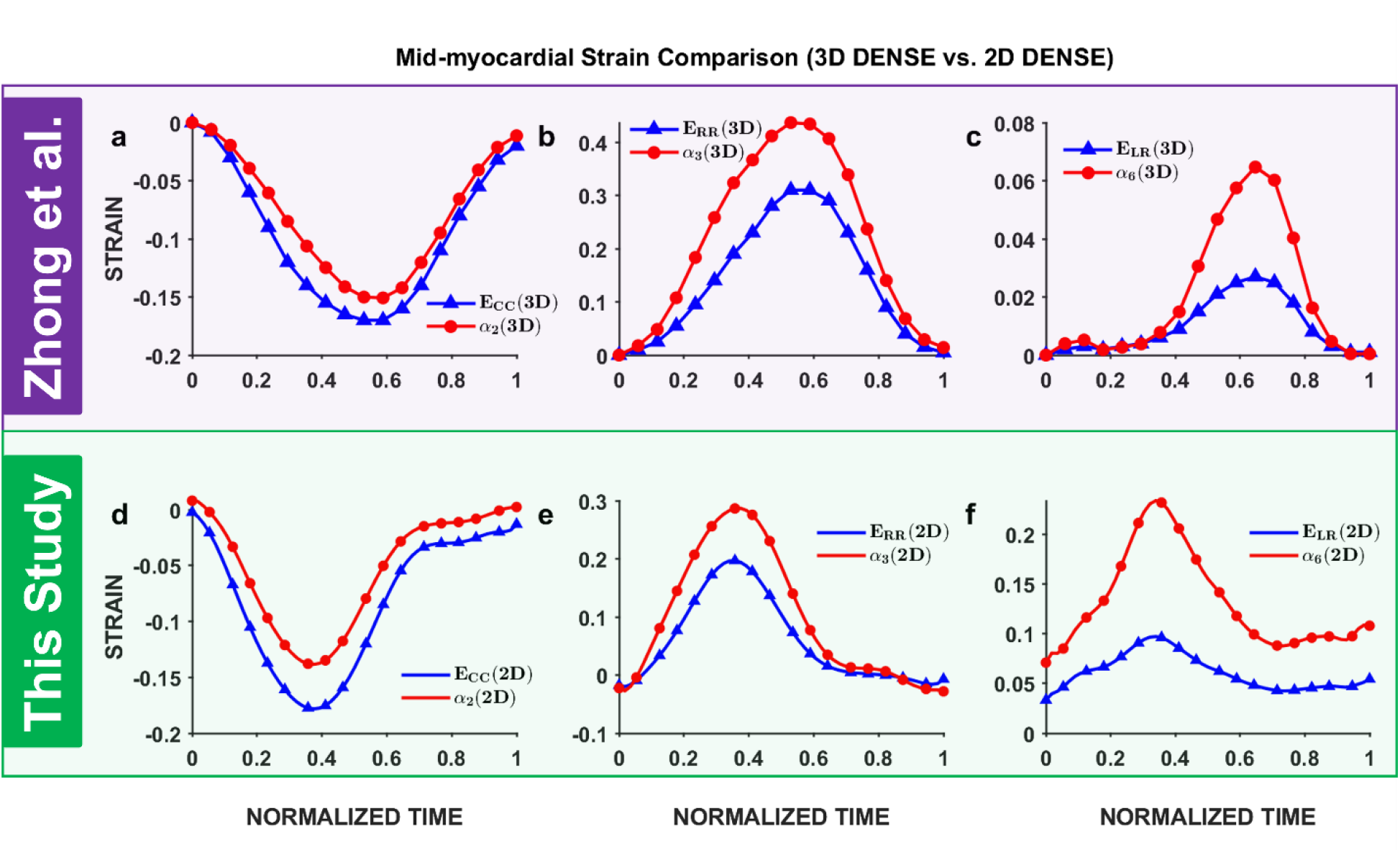
Comparison of the relationship between the mid-myocardial Green strains and their respective strain attributes. The first row shows the Green strains reported in Zhong et al. via 3D DENSE (shown in blue) with their calculated strain attributes (shown in red). The second row shows the mean Green strains obtained in this paper, based on 2D DENSE, (also shown in blue) and their respective strain attributes (shown in red). ECC: circumferential Green strain, E_RR_: radial Green strain, E_LR_: longitudinal-radial Green shear strain, α_2_: mean fiber strain attribute, α_3_: mean laminar thickening strain attribute, and α_6_: crossfiber-radial shear strain attribute. The relationship between the Green strain components and respective strain attribute counterpart from 3D DENSE is at least 87% similar to the relationship between both strain definitions obtained from 2D DENSE.

## DISCUSSION

The goal of this paper was to quantify the 3D deformation of the healthy left ventricular myocardium relative to its mean material directions (fiber, cross-fiber, and radial) using pixel-wise displacement data from DENSE MRI. From the resulting displacement field, an upper triangular stretch tensor was obtained to quantify the normal and shear stretches along the material axes. This stretch tensor was subsequently separated into a dilatation and distortion, and strain was quantified via logarithmic strain attributes. The resulting strains were compared with other widely used definitions of strain, such as the Green strain and engineering strain. The main findings herein were: (1) engineering strain predicts a significantly different deformation pattern of the myocardium compared to that of the Green strain and strain attributes; (2) the logarithmic strain attributes predicts a greater magnitude of deformation per direction than their Green strain counterparts (with the exception of the mean fiber strain attribute *α*_2_); (3) the strain attributes vary from base to apex, and vary regionally within each slice, where the LV lateral wall tends to show greater deformation overall relative to the septal wall; and (4) the myocardium was shown to be compressible, where its change in volume increased from base to apex. Hence, this work sought to establish a more robust method of quantifying myocardial deformation using a strain definition that is more congruent with its mesostructural architecture and better at capturing its physiology. The implications of these findings are detailed below.

### Which strain definition is best suited for the myocardium?

While both the engineering strain and Green strain have been commonly used to quantify myocardial deformation, they fundamentally predict different deformation patterns for most soft tissues. Engineering strain, which quantifies the fractional shortening or elongation, assumes the deformation to be infinitesimal (< 2%). However, it is clear from **Figures 7 – 10** that myocardial deformation far exceeds that into the finite regime. **Figures 7 – 10** further shows how the engineering strain underestimates the deformation when the stretch ratio is greater than 1 (such as in myocardial radial thickening) and overestimates it when the stretch ratio is less than 1 (such as in circumferential or longitudinal shortening), when compared to the Green strain, which is appropriate for finite deformations. Hence, while engineering strain has been used in numerous studies and clinical trials to assess myocardial function, engineering strain should be avoided for modeling the myocardium as it is simply not appropriate [1–4,11,14,26]. Furthermore, engineering strain results from a linearization of the Green strain, i.e., when infinitesimal deformations are assumed (which is an inappropriate assumption for the myocardium), the Green strain simplifies to the engineering strain [61,62].

#### The Polar Green Strains

The Green strain is suitable for “large” or finite deformations; however, careful consideration reveals a few limitations when applied to the myocardium. Primarily, the Green strain is often calculated relative to a global cardiac coordinate system to quantify myocardial deformation. While such strains reveal the geometric changes of the myocardium, it does not explicitly elucidate changes to the mesostructure, such as changes in the myocardial sheetlets and the fibers. Such changes would be more clinically intuitive and congruent with myocardial physiology. Additionally, it is possible that various pathological conditions can lead to similar changes in geometry but have significantly different mesostructural changes. Hence, a mesostructurally informed strain description would be more useful and physiologically intuitive and can serve as a bridge between the macrostructure and the microstructure. Another limitation is that, fundamentally, the Green strain depends on the square of the stretch, which is subsequently subtracted by unity and then halved. This makes its physical interpretability rather difficult, particularly when attempting to use it to describe myocardial changes undergoing various pathophysiological phenomena. Another result of this definition is that the Green strain is “positively biased”. This comes from making **U** equal to **0** in the Green strain equation — indicating that, theoretically, if the stretch was null (which is a physical impossibility as it would imply the material vanishing, thus violating the conservation of mass), the theoretical lower limit of the Green strain is −0.5 (because 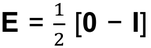). Hence, the range of all possible values are for the Green strain are (−0.5, ꝏ). This could be relevant when considering growth and remodeling adaptations, where the deformation would change due to both motion as well as growth. For example, in certain conditions such as in a myocardial infarction, where the radial thickness of the infarcted zone significantly decreases compared to its baseline, this positive bias would imply that this growth strain is no less than −0.5, even though the thickness of the infarcted zone may have decreased by 60% or above its baseline thickness [63]. This could be a consideration in longitudinal studies when growth adaptations are interpreted relative to a baseline geometry. Finally, the Green strain does not inherently separate dilatation from distortion, which are two separate components of deformation and would provide separate mechanistic insights into the mechanics of the myocardium. If the myocardium was incompressible, such a separation would not be needed, yet as shown herein and in other studies, the compressibility of the myocardium requires such a separation [64–66].

#### The Mesostructural Strain Attributes

To address these limitations in the Green strain, the strain attribute-based definition proposed herein is a logarithmic strain that is derived from material-based stretches. This achieves two main benefits: (1) it yields a physical description of the deformation; and (2) it relates the deformation to changes in the mesostructure. Regarding the physical description, the definition of the logarithmic strain is derived from the fact that the deformation along a certain length is divided into infinitesimal deformations, which are summed together to achieve the net deformation (**Eq. 3**). Hence, logarithmic strain yields the *physical* deformation along the path of interest (unlike the Green strain, which is based on a square of the stretch). Additionally, to describe physical deformation along the material directions, the strain attributes were derived from the upper triangular stretches (which represent the stretches along the material directions). The logarithmic strains that are derived from upper triangular stretches yields a physical description of the deformation that is more intuitive than other strain measures [67]. Therefore, the strain attributes herein are more robust than other strain definitions for describing myocardial deformation.

With a robustly defined strain measure, myocardial deformation can be quantified relative to the mesostructure in a variety of disease and clinical scenarios. For instance, in stem-cell based 3D tissue-engineered constructs (or 3D cardiac patches) aimed at replacing infarcted tissue in myocardial infarction, the deformation of the myocardium via strain attributes before and after implantation can be utilized to measure the degree of restoration of myocardial function (or lack thereof). Additionally, in a variety of clinical trials aimed at testing therapeutics on patient outcomes, mesostructure-based strain attributes can be used as a robust assessment on how these therapeutics can change myocardial function. These strain attributes as markers are not only specific to the myocardium, but also relate directly to the mesostructure, which if changed by the disease, it would be captured by these strain attributes better than other strain definitions.

### Temporally, what do the global strain attributes elucidate about myocardial deformation that the Green strains do not?

#### The Normal Strains

For the global normal deformations, **Figures 7 and 8** show that the mean laminar thickening strain attribute *α*_3_∼ (27 ± 2.3)% is roughly double in magnitude compared to the mean fiber strain attribute *α*_2_ ∼ (−15 ± 0.9)% at end-systole. Physiologically, these values are consistent with studies that found that the individual fibers shorten by ∼10–15% while the LV myocardial wall thickens by ∼30–40% [7,8,15,16,22]. The explanation behind this phenomenon lies in the organization of the myocardial microstructure, where there is more extracellular collagen within the sheets than there is between the sheets (i.e., within the cleavage plains). Additionally, the myofibers contain the protein titin which adds to the overall stiffness [22]. Thus, these coupled effects result in posing greater resistance upon the myocardium to deform along the fiber direction than along either of the other two orthogonal directions (i.e., laminar thickening) [7,8,15,16,22,68]. Thus, these global strain attributes elucidate what the global mesostructural function of the myocardium is. As a result, they have potential in elucidating how this paradigm is changed or impaired in various disease conditions. However, for the geometric Green strains, the magnitudes of E_CC_ and E_RR_ are rather comparable with E_CC_ = (−18 ± 1.7)% and E_RR_ = (22 ± 5.1)%. This difference may be for two reasons: (1) how the Green strain is fundamentally calculated (as explained in the previous section); and (2) these Green strains do not separate the dilatation from distortion. For the second point, such a separation would be immaterial if the myocardium was indeed incompressible; however, as shown in the results and explained in a section below, this is not the case. Even so, the isochoric Green strain improves the calculation of the radial thickening 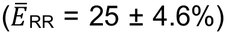 but tends to overestimate the fiber shortening 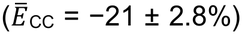, with both strains still relatively comparable. Hence, even though the coordinate system is comparable in this study across all 3 strain definitions (i.e., Green strains, isochoric Green strains, and strain attributes), the strain attributes yield deformations more consistent with the mesostructural physiology of the myocardium.

#### The Shear Strains

For the global shear deformations, the strain attributes reveal greater magnitudes of deformations than their respective Green strain counterparts. Furthermore, *α*_4_ (which quantifies twist) was (4.6 ± 12)%, *α*_5_ (which quantifies laminar imbrication due to longitudinal tilting) was (22 ± 1.2)%, and *α*_6_ (which quantifies laminar imbrication due to circumferential tilting) was (30 ± 7.0)%. While these strain attributes themselves have not been previously quantified for the myocardium, the deformation patterns they seek to describe have been measured in previous studies [5,7,8,15,16,22,35,69–75]. Granted, the studies compared herein were either done by tracking gold-bead markers via biplanar x-rays in large animals or MRI tagging in human subjects, with a few studies that did DENSE [5,7,8,15,16,22,35,69–75]. Hence, the animal studies will be used to qualitatively compare the strain attributes while the human studies will be used for quantitative comparisons of the Green strains. Overall, studies that explicitly sought the measurement of these shear strains in large animals relative to the myocardial microstructure yielded that the cross-fiber-radial shear strain had the greatest magnitude in the myocardium followed by the fiber-radial shear strain (as these two strains contribute to the overall thickening of the myocardium via the imbrication of its sheetlets) [5,7,8,15,16]. The fiber-cross-fiber shear strain (which controls myocardial twist) had the least magnitude [5,7,8,15,16]. The mechanism behind this observation was explained by Dokos et al. as shearing the fibers within sheetlets possesses more cellular and extracellular ‘stiff’ components than shearing the sheetlets along the radial axis or longitudinal axis, requiring more myocardial energy due to being stiffer loading paths [68]. As the stiffness increases, the compliance decreases along that direction, resulting in less motion and thus lower magnitudes of shear strain [68]. While these studies used the Green strain in myofiber coordinates, these strain attributes qualitatively predict myocardial physiological behavior consistent with other similar studies [5,7,8,15,16,22,35,69–75]. Quantitatively, these values are different from some of these studies in human subjects, which will be compared with the Green strains herein, whereby *E*_CL_= 1.3 ± 3.7%, *E*_CR_ = 7.0 ± 0.4%, and *E*_LR_ = 12 ± 2.2%. To illustrate, Kuijer et al. reported shear angles rather than shear strains, but when converted, resulted in greater magnitudes than what was estimated in the work herein, where *E*_CL_∼ 28%, *E*_CR_ ∼ 46%, and *E*_LR_ ∼31%, which does not follow the qualitative comparability in the work herein [71]. This could most likely be due to the limited spatial resolution in tagged MRI, where the resulting uncertainty in the estimating the normal strains can propagate to the shear strain (based on their equation) [71]. On the other hand, other studies have obtained shear strains that are lower in magnitude than the work herein with some of the shear strain components negative in some cases [69,76]. Regarding the sign, while the *E*_CL_ at the base was negative, both *E*_CR_ and *E*_LR_ were positive due to the way they were calculated in this work (cf. **Eqs. 4** and **5**). For the magnitudes, because shear is limited to the number of displacement seed points (indicated by the resolution of the tag lines), MRI tagging often underestimates shear strain. For the analogous DENSE studies, the shear strains herein were more comparable, with slight differences from the Zhong et al. study [35]. Furthermore, the mid-ventricular *E*_CR_ obtained herein is comparable to that of Zhong et al.; however, *E*_LR_ and *E*_CL_ reported herein are slightly greater. This could be because the work of Zhong et al. utilizes 3D DENSE whereas this work takes multiple 2D DENSE slices, map regions based on the AHA bullseye, and assumes uniformity across elements in both size and distribution of strains.

### Regionally, what do the local strain attributes elucidate about myocardial deformation that the Green strains do not?

As shown in **Figure 14**, both *α*_2_ and *α*_3_ show that there is greater fiber shortening and laminar thickening, respectively, on the anterior and lateral walls than those on the septal and inferior walls of the left ventricle. However, *α*_6_ shows that there is greater laminar transverse shearing on the septal and inferior walls than on the anterior and lateral wall. When compared with their Green strain counterparts, *E*_CC_, *E*_RR_ and *E*_LR_ show greater spatial homogeneity. The variation in the strain distribution within the AHA bullseyes in **Figure 14** can be considered segment-to-segment within each short-axis slice (anterior, anterolateral, inferolateral, inferior, inferoseptal, and anteroseptal) as well as slice-to-slice (base, mid, and apex).

#### E_CC_ vs. α_2_

For *α*_2_ and *E*_CC_, their respective bullseye plots in the first two panels of the upper row in **Figure 14** can be compared. As already demonstrated by the bullseye plots in **Figure 14**, *α*_2_ shows greater spatial heterogeneity in the deformation than that of *E*_CC_. Granted, the *p*-value bullseyes show that there were no significant differences between both strain components in the basal segments (except for that of the inferior basal segment). However, careful consideration of how *α*_2_ varies segmentally within each slice demonstrates the mean fiber shortening strain *α*_2_ in the basal and mid slices increases along the anterior and lateral wall segments, decreases along the inferolateral and inferior segments, and then increases again along the inferoseptal and anteroseptal segments. For the apical slice, *α*_2_ increases in the anterior and anterolateral segments, decreases along the inferolateral and inferior segments, and continues to decrease (with a lower gradient) along the septal wall segments. On the other hand, the *E*_CC_ AHA bullseye in **Figure 14** shows relative homogeneity along the anterior and anterolateral segments, an increase along the inferolateral and inferior segments, and relative homogeneity along the inferoseptal and anteroseptal segments. For the slice-to-slice variation, the *α*_2_ AHA bullseye shows that *α*_2_ decreases from base to mid but then increases from mid to apex. However, the *E*_CC_ AHA bullseye shows that *E*_CC_ overall decreases from base to apex for all segments. These results are consistent with that of Moore et al. and Young et al., where they reported the greatest shortening to be at the apex as well as at the anterior and lateral walls [69,76]. Interestingly, for *α*_2_, while **Figure 14** shows segmental heterogeneity throughout each slice, **Figure 11** shows that *α*_2_ averaged over the base, mid, and apex is uniform, which is consistent with previous studies that obtained a uniform regional distribution of the fiber strain [34,44,70].

#### E_RR_ vs. α_3_

For *α*_3_ and *E*_RR_, their respective AHA bullseye plots in the first two panels of the middle row in **Figure 14** can be compared. As already demonstrated by the bullseye plots in **Figure 14**,

*α*_3_ shows greater spatial heterogeneity in the deformation than that of *E*_RR_. Granted, the *p*-value bullseyes show that there were no significant differences between both strain components in all the basal segments (except for the anterolateral and inferoseptal segments), the mid anteroseptal segment, and all the apical segments (except for the inferior segment). However, careful consideration of how *α*_3_ varies segment-to-segment within each slice demonstrates the mean laminar thickening strain *α*_3_ in the basal and mid slices increases along the anterior and lateral wall segments (with a greater gradient in the mid), decreases along the inferolateral and inferior segments, and then increases again along the inferoseptal and anteroseptal segments (but decreases along the anteroseptal segment in the mid slice). For the apical slice, *α*_3_ increases in the anterior and anterolateral segments, continues to increase along the inferolateral and inferior segments (but with a lower gradient), and decreases along the septal wall segments. On the other hand, the *E*_RR_ AHA bullseye in **Figure 14** shows a relatively similar trend but with lower gradients on the anterior and lateral wall segments and higher gradients in the inferior and septal wall segments. For the slice-to-slice changes, the *α*_3_ AHA bullseye shows that *α*_3_ decreases from base to apex for the anterior and lateral wall segments yet increases from base to apex for the inferior and septal wall segments, with the greatest gradient along inferior wall segment. However, the *E*_RR_ AHA bullseye in **Figure 14** shows that *E*_RR_ decreases from base to mid for the anterior and lateral wall segments. The rest of segments show little to no change across the slices (indicating homogeneity from base to apex), with the exception of the inferior wall — with a decrease from base to mid and an increase from mid to apex. These results are consistent with that of Moore et al. and Young et al., where they reported the greatest thickening to be at the anterior and lateral walls [69,76]. However, they reported the greatest thickening near the apex [69,76]. While the thickening magnitudes were great at the apex, they were greatest at the base. It should be noted that in these studies, the myocardium was assumed to be incompressible which would overestimate the thickening magnitudes [69]. As shown in the work herein, this is clearly not the case. Additionally, *α*_3_ is calculated from the ratio of radial thickening to cross-fiber shortening, and while the radial thickening decreases from base to apex, the increasing cross-fiber shortening from base to apex keeps *α*_3_ relatively high throughout the myocardium (compared to *E*_RR_).

#### E_LR_ vs. α_6_

For *α*_6_ and *E*_LR_, their respective AHA bullseye plots in the first two panels of the lower row in **Figure 14** can be compared. As already demonstrated by the bullseye plots in **Figure 14**, *α*6 shows greater spatial heterogeneity in the deformation than that of *E*_LR_. Additionally, the *p*-value bullseyes in **Figure 14** show that the strain patterns for each of the strain measures were significantly different for all segments in the bullseye. Furthermore, the regional magnitudes of *α*_6_ were greater than that of their *E*_LR_ counterparts. For example, the magnitude of *α*6 can reach ∼0.45 in some of the basal segments, yet the magnitude of *E*_LR_ ∼ 0.15 for the same basal segment. Interestingly, within each slice, both the *α*6 and *E*_LR_ bullseye plots demonstrate a qualitatively similar pattern, whereby the strain increases along the anterior and lateral wall segments. These strains decrease along the inferior and inferoseptal segments but increase again along the anteroseptal segment, with the greatest change in the mid slice. The difference between *α*6 and *E*_LR_ is the quantitative magnitudes of their respective segment-to-segment variation, with a greater variation for *α*_6_ given *α*6 is regionally greater than *E*_LR_ in the same regions. Similarly, for variation across the slices, both the *α*6 and *E*_LR_ AHA bullseye plots demonstrate a qualitatively similar yet quantitatively different pattern. Furthermore, in the anterior and lateral wall segments, *α*6 and *E*_LR_ decrease from base to mid but increase from mid to apex, with a greater variation for *α*6. A similar pattern for both *α*6 and *E*_LR_ can be observed for the inferior and septal walls, but with a steeper change from base to mid. Once again, the main difference between both strains is that the variation of *α*6 is greater than that of *E*_LR_. To compare the work herein with similar work of Moore et al., their reported *E*_LR_ in the anterior segment was less than any other segment within the same slice, which indicates an increase at the anterior segment [69]. While this is the case for the anterior wall *E*_LR_ obtained herein, the lateral wall segments would reveal greater magnitudes than that in the anterior segments (in addition to the septal segments in the mid slice). This segmental variation result is similar to what the Young et al. study obtained for *E*_LR_, where the *E*_LR_ in the lateral wall was greater than that in either the anterior or inferior wall [76]. Additionally, the greater septal wall values in the mid slice was also obtained by Kindberg et al [73]. These differences between these studies and the work herein might be due to the use of tagging, where the spatial resolution for shear displacements is less than that of DENSE, which would result in greater uncertainty and thus more prone to error [72]. This might also explain why their *E*_LR_ in some segments were zero, when it has been shown in other studies to be a main contributor to myocardial thickening [5,7,8,68,74]. However, for their variation across slices, they reported greater values of *E*_LR_ in the apex than in the base, which is consistent with the analagous results for *E*_LR_ obtained herein [69]. To explain why *E*_LR_ has lower magnitudes than *α*6, the differences between both strain may perhaps be due to the fundamental difference between both strain measures. For the Green strain, it is obtained from 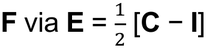, where **C** = **F^T^F** as explained before. For a simple shear experiment (as that conducted in Dokos et al. and Li et al.), where **F** = **I** + γ**e_2_**⊗ **e_3_**, both the shear strain components E_23_ = E_32_ = γ/2 [68,77]. Thus, the Green strain halves the magnitude of shear, yet for the same **F**, the upper triangular **f** = **F**, and thus *α*6 = γ and preserves the magnitude of shear. Hence, the shear strain attributes calculated herein are equal to the amount of shear whereas the shear Green strains halves it, per the definition of each strain.

### Is the myocardium compressible?

One of the main findings of this study is that the myocardium is *not* incompressible (where incompressibility is a common assumption), which is defined as *J* = 1 or *α*1 = 0, shown in **Figures 15 and 16**. Furthermore, the slice-to-slice variation of both *J* and *α*1 show that the compressibility or reduction in myocardial volume at end-systole increases from base to apex. This measurement was enabled using the upper triangular matrix, which made calculating *J* simpler than previous attempts as *J =* det(**F**) = det(**f**) [65,66]. Additionally, the DENSE-measured through-plane stretch *f*_33_ was found to deviate significantly from the *f*_33_ predicted from incompressibility (shown in supplemental materials). While this work might be the first to measure the change in myocardial volume from DENSE-based 3D kinematics using the upper triangular framework and demonstrate that the myocardium is compressible, this idea—that the myocardium is compressible—is not new. Early large animal *ex vivo* and *in vivo* investigations of canine myocardium have shown that during systole, the increased intramyocardial pressure leads to the extrusion of blood into the downstream coronary venules and prevents the inflow from the upstream coronary arterioles [64,78–80]. Other studies have quantified the amount of reduction in blood volume to be 15% – 20% for a myocardial cube of ∼1 cm^3^ and 42% for the LV lateral wall [64,80–82]. More recent *in vivo* investigations in sheep have shown a reduction by 7% at the basal anterior wall and 10% at the midwall while another investigation has shown an end-systolic *J* of 0.85 with an increasing *y*-gradient from base to apex [66,83,84]. Other *in vivo* studies in humans from MRI found that the ratio of the systolic to diastolic myocardial volume (calculated from the whole LV myocardium) was ∼0.87 [65]. Additionally, Avazmohammadi et al. calculated a 16% reduction in volume from the 3D DENSE strains measured by Zhong et al. [35,66].

*Spatial Distribution of J and α1*

The regional distribution of *J* and *α*1 is shown in their respective AHA bullseye plots in **Figure 15**. As already demonstrated by the bullseye plots in **Figure 15**, both *J* and *α*1 show relative homogeneity in the myocardial compressibility with the exception of a few segments, such as that of the basal and mid septal segments. Furthermore, from **Figure 15**, *J* can reach a magnitude of ∼ 0.80 – 0.90 near the base (corresponding to an *α*1 range of ∼ − 0.10 to − 0.22) and 0.75 – 0.80 near the apex (corresponding to an *α*1 range of ∼ − 0.22 to − 0.30). To get a better physical understanding of that these values mean, a *J =* 0.90 would, for example, indicate that the end-systolic myocardial volume is 90% of its reference volume at end-diastole (i.e., a 10% reduction in volume). The temporal evolution of the change in myocardial volume is also shown in **Figure 15**. From the bullseyes in **Figure 15**, both the *α*1 and *J* bullseye plots demonstrate a relative segment-to-segment homogeneity across each short-axis slice, except at the basal and mid inferoseptal segments where there is an increase, indicating a greater *J* or a less reduction in myocardial volume. Thus, along each short-axis slice, the segment-to-segment volume change is consistently greater on the anterior, lateral, and inferior walls yet lower in the septal wall segments. However, for the slice-to-slice variation, both the *α*1 and *J* bullseye plots show a decrease from base to apex, indicating a greater reduction in volume along the base-to-apex axis, which is also shown in the decreasing *J* values (or increasing *α*1 values in **Figure 16**). This is also indicated by increasing magnitudes of volume strains (i.e., *α*1) in from base to apex, both demonstrated in **Figures 15 and 16**.

#### Physiological Drivers of Myocardial Volume Reduction

Physiologically, most of the values for the reduction in volume obtained herein can be explained by a reduction in the vascular volume (which is reported to be roughly 15% – 20%) [64,66,82]. However, higher values for *α*1 such as that in the apex and certain segments in the mid slice suggest that there are perhaps other additional mechanisms at work. To illustrate, Ashikaga et al. proposed three possible mechanisms: (1) potential anatomical communications between the coronary vessels and the ventricular lumen (i.e., vascular-luminal communications) where pre-capillary arterioles communicate with the spaces found between the LV trabeculae; (2) the presence of trabecular spaces, predominantly found to extensively line the endocardial layer of the apical region, provide ‘shrinkable’ spaces during systole – which could potentially explain the higher magnitudes of *α*1 near the apex; and (3) changes occurring in the myocardial inter-sheetlet spaces due to sheetlet shear during the cardiac cycle can change the blood volume within [85]. The increase of *α*6 along the mid and apical slices in **Figure 15** could also potentially explain the greater values *α*1 in these regions. Hence, this observation would also imply that myocardial distortion and dilatation, while mathematically separable, are mechanistically coupled.

#### Myocardial Compressibility vs. Squeezability

Finally, an important concept that must be acknowledged herein is the definition of compressibility. Conventionally, compressibility is defined as the ratio between the change of volume in the deformed configuration (*∂v*) to the change of volume in the undeformed configuration (*∂V*). Using the relationship between mass density *ρ*, mass *m*, and volume *v* (i.e., *m* = *ρV*), *J* can be rewritten in terms of the density of the deformed configuration *ρ* and that of the undeformed configuration *ρ*_0_, such that *J = ∂v*/*∂V* = *∂ρ* /*∂ρ_0_*. Hence, an important assumption for *J* that seems to be overlooked is that mass is conserved — a defining characteristic for a closed system. This implies that for a compressible material, the *same mass* occupies a smaller volume. However, as proposed by the aforementioned mechanisms of myocardial compressibility (with the exception of the reduction of the inter-sheetlet spaces due to myocardial shear), most of this change in volume can be attributed to blood extruded ‘out’ of the myocardium, and into either the coronary sinus through its Thebesian drainage or the ventricular lumen via the arterio-luminal communications [85]. It is, therefore, possible that there is loss of myocardial mass from end-diastole (its undeformed configuration) to end-systole (its deformed configuration). It is also important to point out that the region of interest in the work herein was considered over the full thickness of the myocardium, where such vascular effects would be significant. Hence, the conclusion herein is that the myocardium is not compressible per se, but rather ‘squeezable’ where it can extrude some of its mass via a separation of phases (the liquid phase from the solid phase). Here, the liquid phase is blood while the solid phase is the cellular and extracellular components. When the blood mass is subtracted out, the remaining solid phase would be incompressible. This is perhaps why the assumption of incompressibility remains valid for *ex vivo* mechanical testing on both myocardial slabs and myocardial cubes as such specimens are usually devoid of blood flow and obtained from arrested hearts [68,77,86,87]. Granted, incompressibility is often assumed for slabs as it is rather difficult to measure the deformation along the thickness of the slab and it simplifies the math. Even so, *J* has been measured for myocardial cubes that span the myocardial thickness and was found that the incompressibility assumption is valid in such experiments [8]. Additionally, if the myocardium was truly compressible, the volume transition from end-systole to end-diastole would rather be steeper, such as memory foam. Instead, the transition is rather gradual as it is driven by blood flow – the time needed for arterial blood to perfuse the capillary beds during diastole or for blood to drain from the capillary beds into the venous side during systole.

### What are the limitations of this study?

The novelty of this study lies in measuring logarithmic strain attributes along the material directions, but there are limitations. First, to obtain the 3D strains, segments from the short-axis slices were mapped to their respective segments in the long-axis slices. While this method combines these 2D segments to achieve the 3D deformations, it possible that the short-axis slices from different subjects corresponded to slightly different locations along the long axis. As a result, the 3D segmental volume may not have been precisely consistent among all the subjects, but such variations were not significant as the standard deviations did not vary significantly from the mean strains. Additionally, it was the mean strain within each segment that was sought after in this work, which would limit the corresponding uncertainty. However, the deviations would be much more significant if the strains were calculated for each voxel, necessitating the use of registration algorithms for this task.

Second, to find the material-based strain attributes, the mean fiber direction was assumed to be along the hoop direction. While this is generally true, it is not quite accurate. Other studies have reported that while this is case for the basal and mid-myocardial regions, the hoop-aligned fibers occur in the inner one-third of the apex rather than the middle [5,7,8]. Third, the transmural variation of strain across the myocardium was not calculated, which would be an important assessment. To address both limitations, knowledge of the local voxel-wise myofiber orientation would be required, which could either be assumed based on the rule-based algorithm or measured from diffusion tensor imaging (DTI) [88,89]. Because *in vivo* cardiac DTI sequences currently exist, they can be used to measure the voxel-wise fiber orientation and registered with the voxel-wise DENSE displacements to calculate the strain attributes [6,20,21,34,44,89]. Hence, the strain attributes would perhaps be the most suitable and compatible strain measure for registration with DTI.

Finally, since these strain attributes have not been measured before *in vivo*, their values were only qualitatively validated against fiber-based Green strains while their quantitative validation was indirectly assessed through validating their respective polar Green strains from which they were derived. As a result, more studies that would employ *in vivo* 3D kinematics and measurement of these strain attributes would need to be implemented. Other sequences and imaging modalities can be used to quantify these strain attributes, but there would be penalties on the reproducibility of their measured values due to dependence on the spatial resolution of the imaging modality used. Additionally, because cardiac cine imaging is ECG-gated, quantifying myocardial strain would prove difficult in patients with arrhythmias as well as in images with poor quality.

## CONCLUSION

This paper developed strain attributes which reflect the myocardial mesostructure and compared their values to the Green strain, which quantifies the changes in myocardial geometry. The results herein show that (1) the strain attributes were closer at capturing the regional changes undergoing the myocardial sheetlets and fibers than the Green strain; (2) the myocardium is compressible; and (3) engineering strain is inappropriate for the myocardium. The six strain attributes can succinctly describe the 3D deformation of the myocardium as well as connect changes in myocardial physiology to the mesostructure. Capturing 3D deformation along the myocardial material directions allows quantifying microstructural changes, potentially providing new clinical intuition. Measurement of such strains could provide clinicians with more mechanistic insights into the LV remodeling process and may serve as predictive markers of adverse remodeling in various myocardial diseases, which could be a potential consideration to investigate in future clinical studies with patient cohorts. Finally, because these strain attributes can capture the mesostructural deformation, they can be registered with diffusion tensor imaging (DTI) to obtain the voxel-wise deformation of the myocardium relative to its material axes.

## ACKNOWLEDGMENT

The authors would like to acknowledge John C. Criscione, from Texas A&M University, for discussions on myocardial dynamics.

## NOMENCLATURE

|  |  |
|---|---|
| DENSE | Displacement ENcoding with Stimulated Echoes |
| *J* | Incompressibility Constraint (Jacobian) |
| *α*<sub>1</sub> | Mean Dilatational Strain |
| *α*<sub>2</sub> | Mean Fiber Strain |
| *α*<sub>3</sub> | Mean Laminar Thickening Strain |
| *α*<sub>4</sub> | Mean Laminar Fiber–Cross-fiber Shear Strain (Shear due to Laminar Twist) |
| *α*<sub>5</sub> | Mean Laminar Fiber–Radial Shear Strain (Shear due to Laminar Tilting about the Circumferential Axis) |
| *α*<sub>6</sub> | Mean Laminar Fiber–Radial Shear Strain (Shear due to Laminar Tilting about the Circumferential Axis) |
| *E*<sub>CC</sub> | Circumferential Green Strain |
| *E*<sub>LL</sub> | Longitudinal Green Strain |
| *E*<sub>RR</sub> | Radial Green Strain |
| *E*<sub>CL</sub> | Circumferential-Longitudinal Green Shear Strain |
| *E*<sub>CR</sub> | Circumferential-Radial Green Shear Strain |

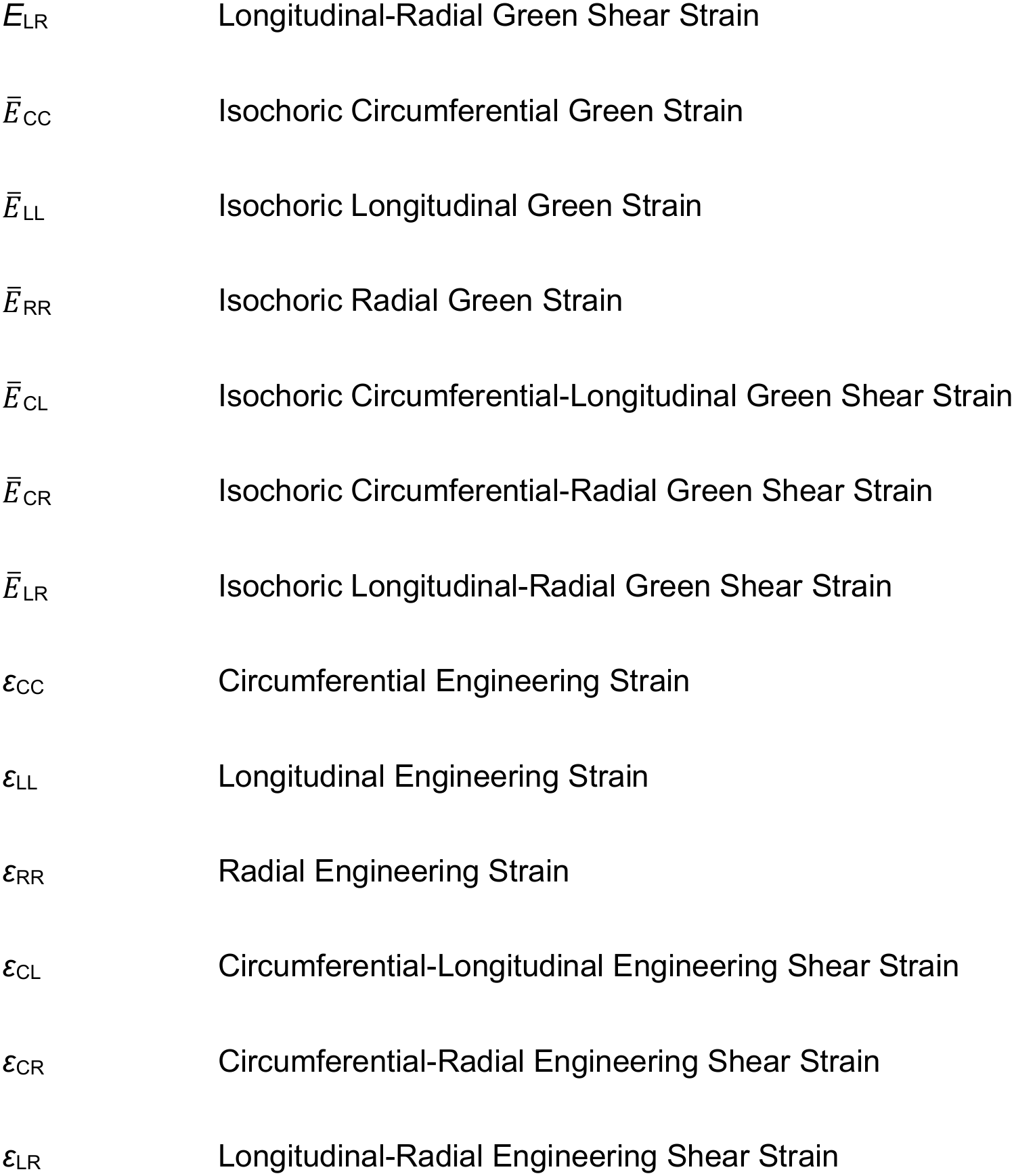

|  |  |
|---|---|
| *E*<sub>LR</sub> | Longitudinal-Radial Green Shear Strain |
| *Ē*<sub>CC</sub> | Isochoric Circumferential Green Strain |
| *Ē*<sub>LL</sub> | Isochoric Longitudinal Green Strain |
| *Ē*<sub>RR</sub> | Isochoric Radial Green Strain |
| *Ē*<sub>CL</sub> | Isochoric Circumferential-Longitudinal Green Shear Strain |
| *Ē*<sub>CR</sub> | Isochoric Circumferential-Radial Green Shear Strain |
| *Ē*<sub>LR</sub> | Isochoric Longitudinal-Radial Green Shear Strain |
| ε<sub>CC</sub> | Circumferential Engineering Strain |
| ε<sub>LL</sub> | Longitudinal Engineering Strain |
| ε<sub>RR</sub> | Radial Engineering Strain |
| ε<sub>CL</sub> | Circumferential-Longitudinal Engineering Shear Strain |
| ε<sub>CR</sub> | Circumferential-Radial Engineering Shear Strain |
| ε<sub>LR</sub> | Longitudinal-Radial Engineering Shear Strain |

