## Supplemental Figures for "Quantification of Local Mesostructural Deformation of the Myocardium using DENSE MRI and a Novel Set of Strain Attributes"

**John N. Oshinski**
Department of Radiology and Imaging Sciences
Emory University School of Medicine
Atlanta, GA, 30332

Wallace H. Coulter Department of Biomedical Engineering
Georgia Institute of Technology & Emory University School of Medicine
Atlanta, GA 30332

### SUPPLEMENTAL MATERIALS

#### Longitudinal Green Strain ($E_{LL}$) vs. Mean Laminar Thickening Strain Attribute ($\alpha_3$)

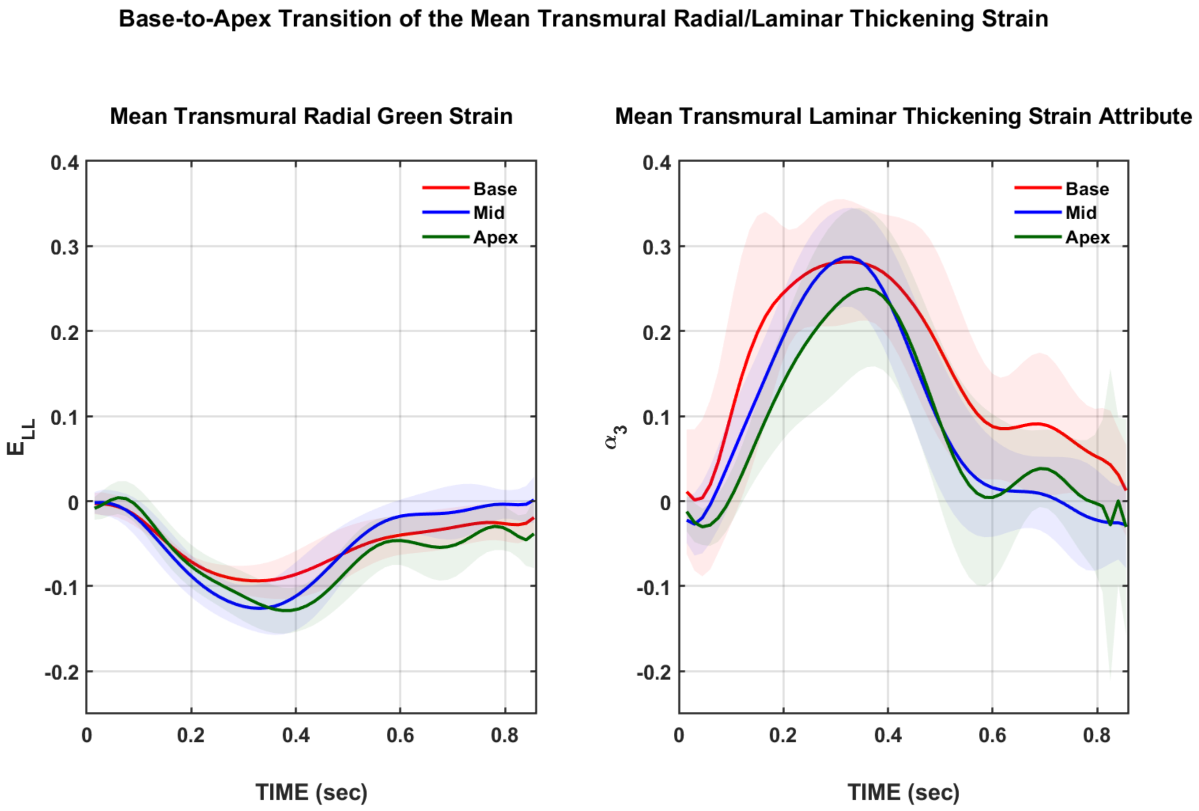


***Figure S1.*** *Base-to-apex transition of the strain in the LV myocardium across a normalized cardiac cycle. The panels show the longitudinal Green strain (left) and the mean laminar thickening strain attribute (right). The curves displayed pertain to the basal (red), mid (blue), and the apical (green) myocardium.*

**Circumferential–Longitudinal Shear Green Strain ($E_{CL}$) vs. Mean Fiber–Cross-fiber Strain Attribute ($\alpha_4$)**

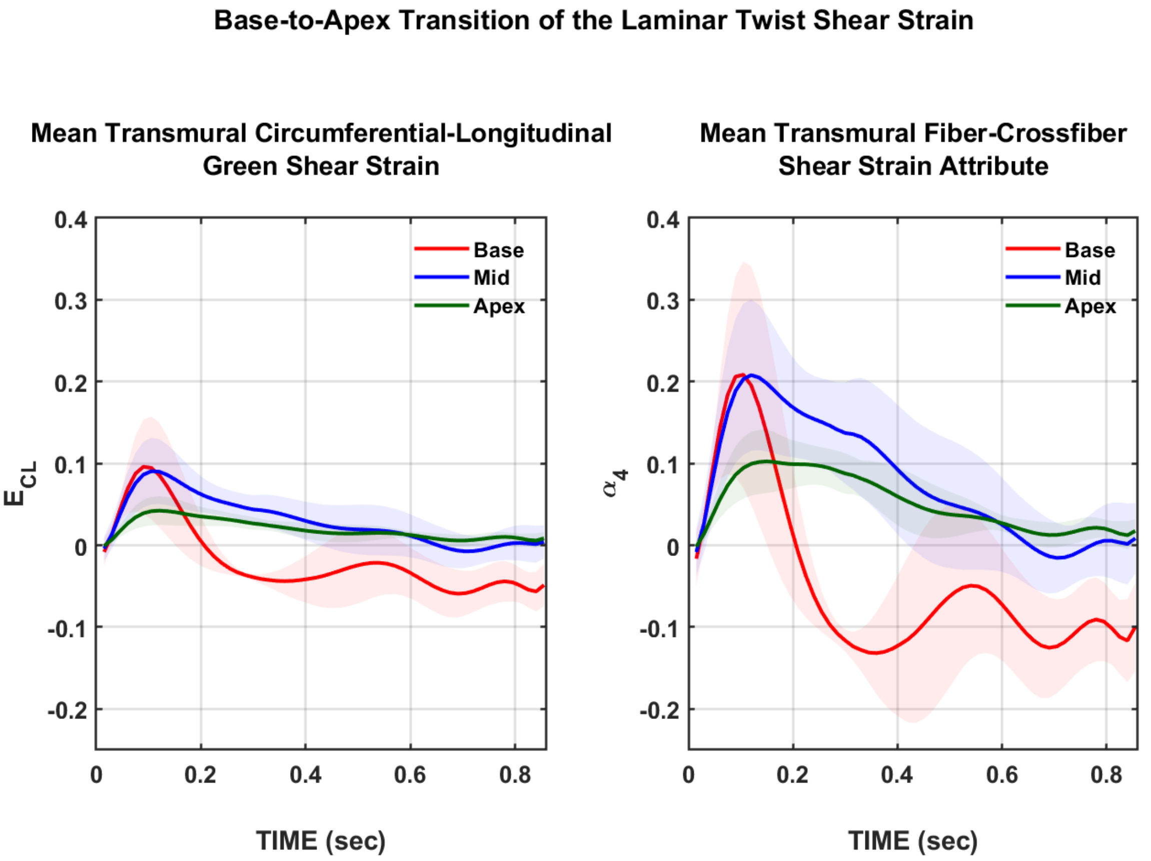


***Figure S2.*** *Base-to-apex transition of the strain in the LV myocardium across a normalized cardiac cycle. The panels show the circumferential–longitudinal shear Green strain (left) and the mean fiber– cross-fiber strain attribute (right). The curves displayed pertain to the basal (red), mid (blue), and the apical (green) myocardium.*

**Circumferential–Longitudinal Shear Green Strain ($E_{CR}$) vs. Mean Fiber–Cross-fiber Strain Attribute ($\alpha_5$)**

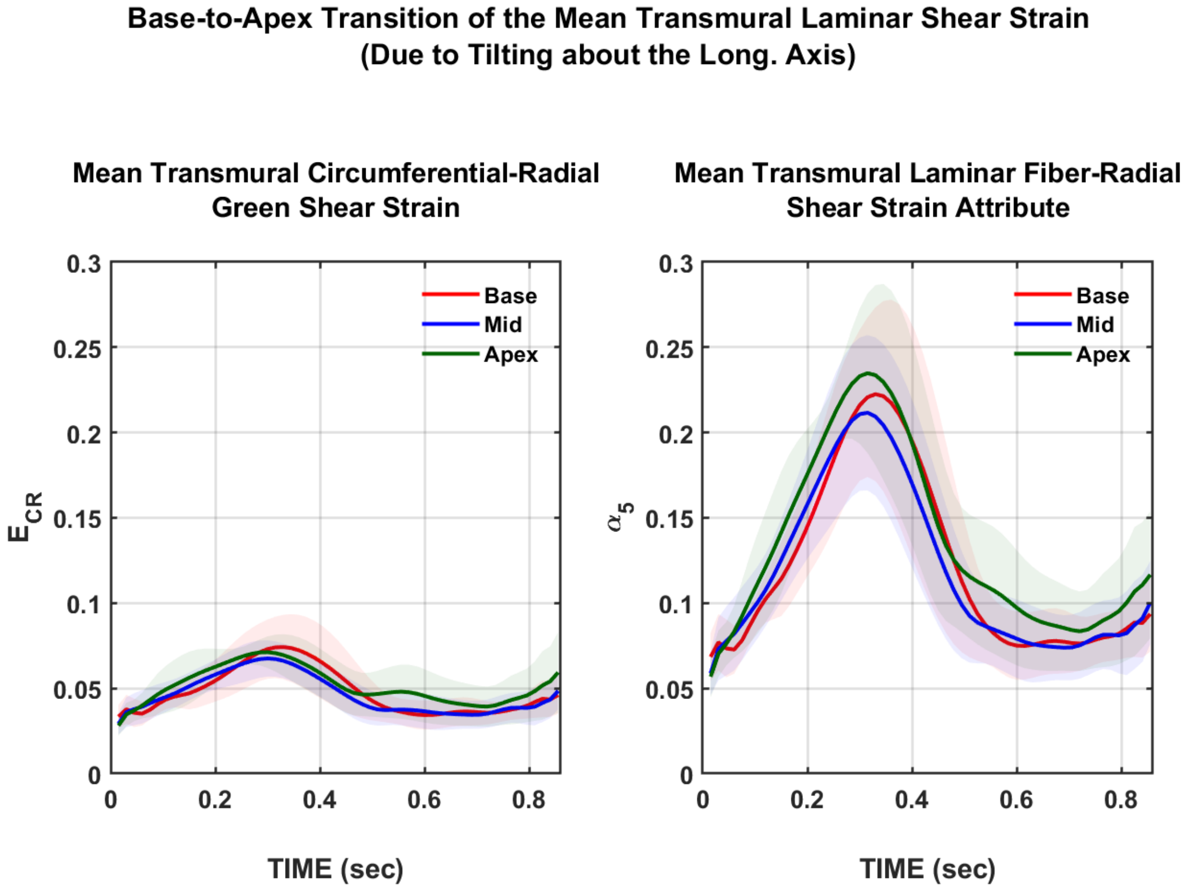


***Figure S3.*** *Base-to-apex transition of the strain in the LV myocardium across a normalized cardiac cycle. The panels show the circumferential–radial shear Green strain (left) and the mean fiber–radial strain attribute (right). The curves displayed pertain to the basal (red), mid (blue), and the apical (green) myocardium.*

### AHA 17-Segment Model of the End-systolic Strains

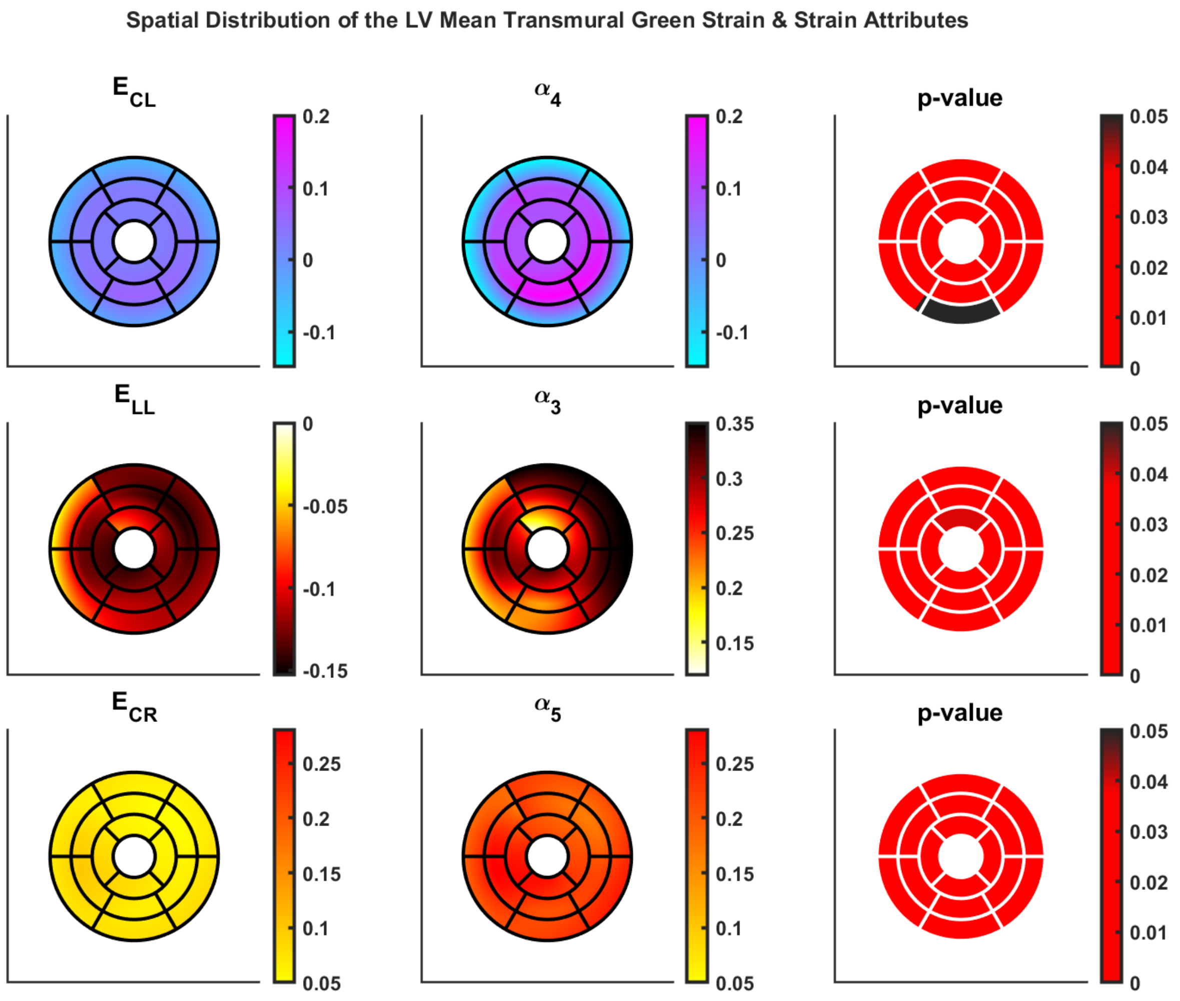


***Figure S4.*** *The spatial distribution of select strain components across the AHA 17-segment model in the LV myocardium with the p-values from a segment-to-segment paired t-test shown on the far right panel. Black segments indicate no significance ($p > 0.05$) while red segments indicate differences that were significant ($p < 0.05$).*

### The Compressibility of the Myocardium

*Measured Through-Plane Stretch vs. Predicted Through-Plane Stretch from Incompressibility*

The measured 3 upper triangular normal stretches (i.e., stretches along the mean material directions – $f_{11}$, $f_{22}$, and $f_{33}$) were plotted over the cardiac cycle in each short-axis slice (i.e., basal, mid, and apical myocardium), displayed below in **Figure S5**. Additionally, the short-axis through-plane stretch predicted from incompressibility (i.e., $f_{22}^{pred} = 1/[f_{11}f_{33}]$), was plotted (shown by the dashed green curve in **Figure S5**). The significant deviation of $f_{22}^{pred}$ from $f_{22}$ can be clearly observed in **Figure S5**. This implies that modeling the ventricular myocardium as an incompressible material would yield significant deviations from its measured behavior.

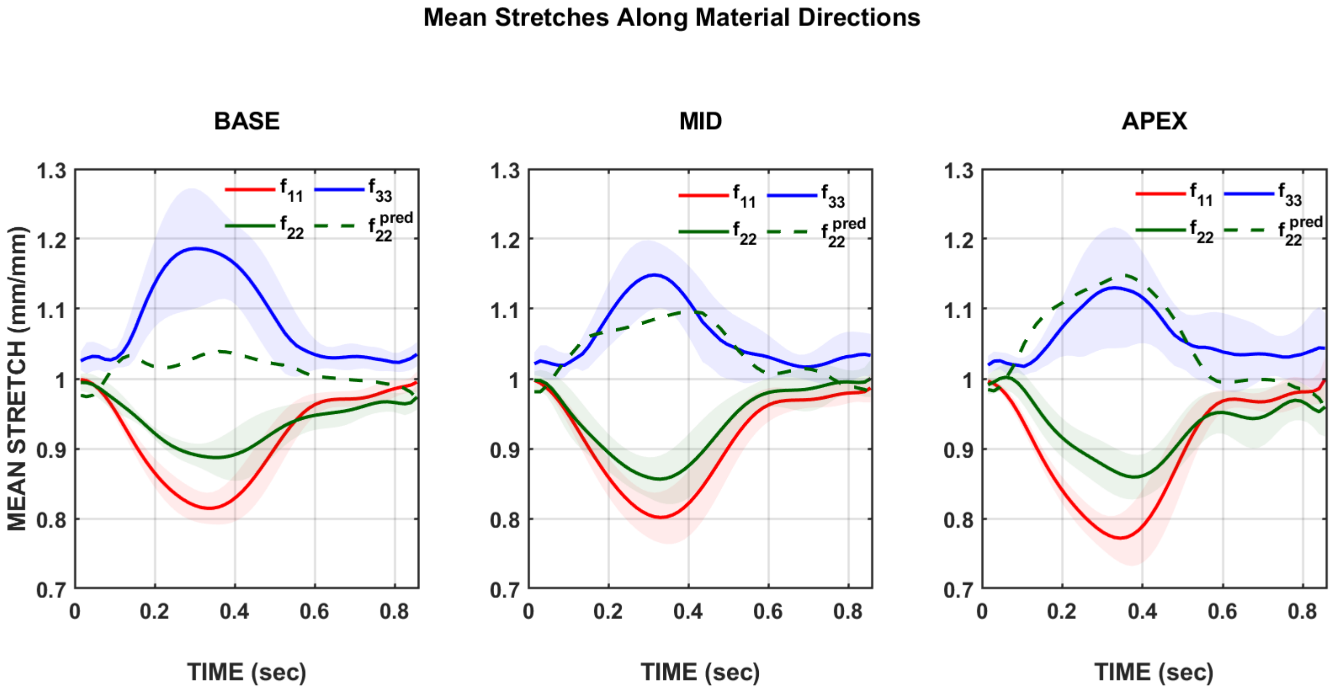


***Figure S5.*** *The temporal variation of the material normal stretches in the basal (left), mid (middle), and apical (right) myocardium over the cardiac cycle. The stretches shown are $f_{11}$ (red), $f_{22}$ (blue), and $f_{33}$ (green).*
